# Temperature–Energy-space Sampling Molecular Dynamics: Deterministic, Iteration-free, and Single-replica Method utilizing Continuous Temperature System

**DOI:** 10.1101/760918

**Authors:** Ikuo Fukuda, Kei Moritsugu

## Abstract

We developed coupled Nosé–Hoover (NH) molecular dynamics equations of motion (EOM), wherein the heat-bath temperature for the physical system (PS) fluctuates according to an arbitrary predetermined weight. The coupled NH is defined by suitably jointing the NH EOM of the PS and the NH EOM of the temperature system (TS), where the inverse heat-bath temperature *β* is a dynamical variable. In this study, we define a method to determine the effective weight for enhanced sampling of the PS states. The method, based on ergodic theory, is reliable, and eliminates the need for time-consuming iterative procedures and resource-consuming replica systems. The resulting TS potential in a two dimensional (*β, ϵ*)-space forms a *valley*, and the potential minimum path forms a *river* flowing through the valley. *β* oscillates around the potential minima for each energy *ϵ*, and the motion of *β* derives a motion of *ϵ* and receives the *ϵ*’s feedback, which leads to a *mutual boost* effect. Thus, it also provides a specific dynamical mechanism to explain the features of enhanced sampling such that the temperature-space “random walk” enhances the energy-space “random walk.” Surprisingly, these mutual dynamics between *β* and *ϵ* naturally arise from the static probability theory formalism of double density dynamics that was previously developed, where the Liouville equation with an arbitrarily given probability density function is the fundamental polestar. Numerical examples using a model system and an explicitly solvated protein system verify the reliability, simplicity, and superiority of the method.

## I. INTRODUCTION

Molecular simulation has become a useful tool for condensed matter, soft-matter, chemical, and biological physics to investigate the characteristics of physical systems (PSs) in terms of microscopic descriptions based on atomic models and their interactions [1–3]. Macroscopic quantities and thermodynamic variables can be obtained by suitably averaging the corresponding phase-space function values that are sequentially generated in the simulation. These outputs are obtained by solving a certain equations of motion (EOM) of the molecular dynamics (MD) method or by a suitable probabilistic procedure in the Monte-Carlo (MC) iteration method [4].

However, the intrinsic problem of these methods is that a long simulation period is necessary to study the macroscopic behavior of the target PS. The first reason for this necessity is the large difference between the unit step of the method and the time required for observing the physical behavior [5]. The former is approximately 1 fs in MD while the latter often goes beyond 1 s. Many efforts have been made to address this problem via computer hardware development [6] and algorithmic techniques such as coarse-grained description [7] and Markov state models [8]. The second reason why these methods take a long time is the slowness underlain in the algorithm, which is slow time development in MD and slow iteration refreshment in MC. This is mainly due to the fact that PS coordinate *x* ≡ (*x*_1_, …, *x*_*n*_) should explore in the complicated surface of the potential energy function *U* (*x*) defined by the interactions of atoms. The surface should have very high barriers, narrow escaping routes, and critical points of various indexes.

Generalized-ensemble enhanced sampling methods have been developed to address these slow development problems: multicanonical molecular dynamics (McMD) [9, 10] and replica exchange molecular dynamics (REMD) [11, 12] are frequently used by many researchers. McMD deforms the energy surface so that the probability of each energy value within a given range is equal. To realize this, *U* (*x*) is redefined iteratively up to the tolerance, where the intractable energy density of state should be handled. REMD is basically a parallel algorithm, where a sufficient number of *N*_R_ replicas of the original PS are prepared at different *N*_R_ temperatures. Individual replica *a* is equilibrated under the Boltzmann-Gibbs (BG) distribution at temperature *T*_*a*_ within a certain period, and the temperature is exchanged among the replicas at a time after the period. The total system should obey the product distribution of these *N*_R_ distinct-temperature BG distributions. These two MD methods are based on MC methods [13–15]. See reviews [16–18] for other methods, various applications, and related topics. Recent developments can be founded in literature [19–24]. The coupled Nosé–Hoover equation (cNH) proposed by the authors [25, 26] also brings many temperatures to the target PS. While NH EOM [27, 28] generates the density ∝ exp(−*βE*(*x, p*)), where *E*(*x, p*) = *K*(*p*)+*U* (*x*) is the total energy of the PS and *β* = 1*/k*_B_*T* is the inverse temperature of the heat bath surrounding the PS, the cNH generates the density *ρ*(*x, p, β*, …) ∝ exp(−*βE*(*x, p*)) *f* (*β*), where *f* (*β*) is an arbitrarily predetermined weight of *β*. This means that *β* is not a constant parameter in the cNH but a continuous dynamical variable that obeys the weight *f* (*β*), and that exp(−*βE*(*x, p*)) reads the density of a conditional probability such that the PS variable (*x, p*) emerges when the inverse temperature takes the value of *β*. The cNH, which consists of the NH EOM of the PS and the NH EOM of the “temperature system” (TS), suitably couples these two EOMs to realize the density *ρ*(*x, p, β*, …). Provided with infinitely many temperatures from the TS, the PS obeys the marginal distribution density

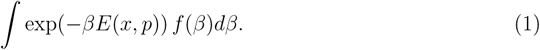

This implies that the integral of uncountably many BG distributions is generated in the cNH, which leads to the potential ability of enhanced sampling. In contrast to the REMD, the cNH only needs the target PS along with the TS, which is very small (typically 1 degree-of-freedom), and the temperature (ex)change is automatically done; the time development of the PS and TS is described by a uniform ordinary differential equation (ODE). Specifically, the cNH brings uncountably many temperatures to the PS automatically via continuous time development using one replica.

In order to make full use of the cNH for enhanced sampling, it is important to know the distribution of *β*, which is

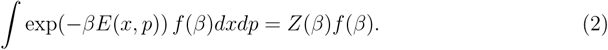

However, the partition function *Z*(*β*) is unknown *a priori* (except for solvable systems) and becomes intractable with increasing system size. Thus, the point is to provide a method to determine *Z* at the required accuracy to be able to define the effectual *f* that leads to give accurate macroscopic quantities via the long-time average. This work provides a solution to this non trivial problem by obtaining sufficient information on *Z* with less iterative procedures and by taking it simply into *f*. We are thus free from heavy tasks, such as those in the McMD method to determine the energy density of state.

The current study clarifies that the cNH accepts the ability of enhanced sampling, not only for the *K*(*p*)-space via Eq. (1), but also for the *U* (*x*)-space; viz., the cNH potentially involves the function of the McMD. We have found its dynamical origin, which leads to a clear statement on why temperature-space sampling leads to effective conformational sampling. In the cNH, the coordinate of the TS, *Q*, which corresponds 1-1 to *β* such that *β* = *σ*(*Q*), moves according to the cNH EOM. The motion in the TS obeys the *TS potential*, which is described by *f* (*β*), the PS energy *ϵ* = *E*(*x, p*), and related variables. By taking the information of *Z* into *f* by the current protocol, the TS potential projected on the two-dimensional (2D) (*β, ϵ*)-space forms a *valley* and takes its minimum nearly along a curve

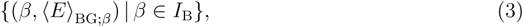

where ⟨*E*⟩_BG;*β*_ is the total-energy average in the BG distribution ∝ exp(−*βE*(*x, p*))*dxdp*. This implies that the movement of *β* triggers the movement of *ϵ* along this curve, and that the enhancement of *β* sampling in the arbitrarily given range *I*_B_ ≡]*β*_min_, *β*_max_[leads to enhancement of energy sampling (see Fig. 1). In other words, the TS in the cNH plays the role of a guide for energy/conformation sampling of the PS by providing a *route* in the (*β, ϵ*)-space. Thus, PS state sampling is not closed within the PS, in contrast to the McMD, but continuously activated by the TS, thus solving the slowness problem in molecular simulation algorithms.

**FIG. 1.**
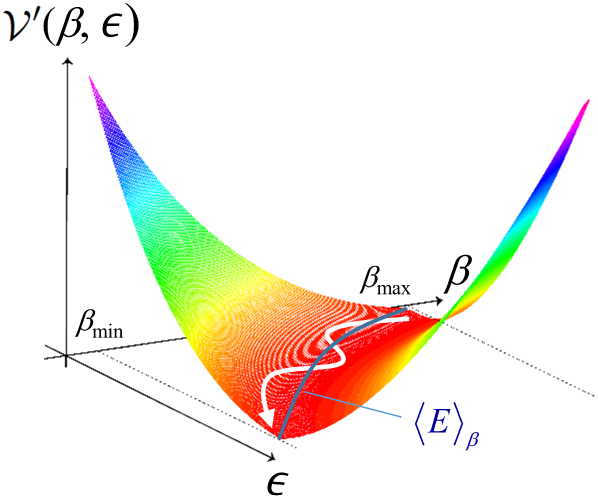
Schematic figure of the TS potential defined in the 2D (*β, ϵ*)-space. The dynamics of the cNH are explored on this surface.

It should be noted that these schemes and their results are not based on a heuristic approach, but on rigorous mathematics and extensible context. The realization of the distribution density exp(−*βE*(*x, p*)) *f*(*β*) in the cNH comes from the existence of an absolutely continuous invariant measure *ρ*(*ω*)*dω* for a continuous dynamical system defined by the cNH vector field. This fact enables us to get any target distribution (e.g., the BG distribution at a target temperature) after enhanced sampling by the cNH via reweighting techniques. It also means that the method does not require any specific reaction coordinate, which is not given *a priori* and often becomes a nontrivial task for conducting methods such as metadynamics [29] and milestoning [30]. The extensibility of the cNH comes from the general framework of double density dynamics [31]. It does not limit to the form of exp(−*βE*(*x, p*)) *f* (*β*), which can be extended to a general form, as will be demonstrated.

In the current work, we present a protocol to easily realize the desired distribution of *β* and sample a wide temperature region. We demonstrate the resulting characteristic sampling features of the PS. Section II reviews the cNH to help readers understand the current work. Section III clarifies our motivation, provides an outline of the method, and introduces its characteristic features. Section IV demonstrates the general framework and technical details of the protocol for constructing an effectual *f*. This is done by giving an approximation of *Z* via a simple initial guess (Sec. IV B) and its refinement (Sec. IV C), along with a smoothing technique to describe *f* for stable use in MD. As detailed in Sec. V, the resulting *f* determines the TS potential, which forms the highlight of the current method and characterizes the PS state sampling. Section VI describes the application of the method to a model system and an explicitly solvated protein system for validation by verifying the marginal distributions, ergodicity, and free energy landscape through comparison with a conventional method. We find that the method works very well in these systems and observe the (*β,ϵ*)-space sampling behavior to confirm its dynamical origins; the discussions conclude in Sec. VII.

## II. REVIEW OF THE COUPLED NOSÉ-HOOVER EQUATION

### A. Equations of motion

We begin by reviewing the cNH EOM [25, 26], represented as

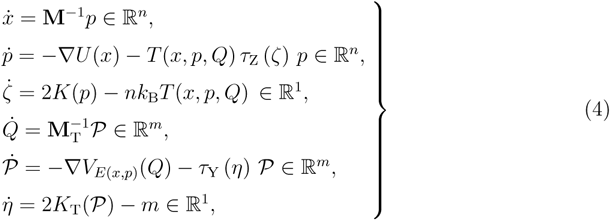

where *x* ≡ (*x*_1_, …, *x*_*n*_) *∈* ℝ^*n*^ are the coordinates for the PS with *n* degrees-of-freedom; *p* ≡ (*p*_1_, …, *p*_*n*_) *∈* ℝ^*n*^ are the corresponding momenta; **M** is masses; *U* (*x*) is the potential energy defined on a domain *D* ⊂ ℝ^*n*^; and 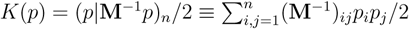 is the kinetic energy, whose instantaneous value is considered proportional to the PS temperature *T*_P_ ≡ 2*K*(*p*)*/nk*_B_. *ζ ∈* ℝ is the dynamical friction variable that controls the temperature of the PS. The variables (*x, p, ζ*) hence form analogue of the conventional NH EOM [27, 28]. See Refs. [32–34] for details on thermostat methods. The main difference between the conventional NH EOM and the first three equations is that the heat bath temperature is not a constant value but a dynamical variable *T* (*x, p, Q*). The *dynamical temperature T* (*x, p, Q*) is controlled by a new (dimensionless) variable *Q* ≡ (*Q*_1_, …, *Q*_*m*_) *∈* ℝ^*m*^, which plays the role of coordinates for the TS. The TS is described by *Q*, corresponding momenta *𝒫* ≡ (*𝒫*_1_, …, *𝒫*_*m*_) *∈* ℝ^*m*^, and an additional control variable *η ∈* ℝ^1^, which is an analogue of *ζ*. Specifically, (*Q, 𝒫, η*) forms an alternative NH EOM, with *V*_*E*(*x,p*)_(*Q*) and *K*_T_(*𝒫*) as the potential and kinetic energies in the TS, respectively. Here, *V*_*E*(*x,p*)_(*Q*) also depends on the PS via the PS energy *E*(*x, p*) ≡ *U* (*x*) + *K*(*p*), and −∇*V*_*E*(*x,p*)_(*Q*) is the force acting on *Q*. The TS kinetic energy is defined by 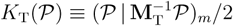 with **M**_T_ representing the *masses* for the TS.

Hence, the phase-space point in the cNH is denoted by *ω* = (*x, p, ζ, Q, 𝒫, η*), with the total phase space Ω contained in ℝ^2(*n*+*m*+1)^ = ℝ^*n*^ × ℝ^*n*^ × ℝ × ℝ^*m*^ × ℝ^*m*^ × ℝ. The cNH EOM is thus described as the ODE, 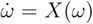, defined on Ω and designed to meet the Liouville equation

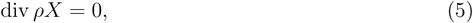

where *ρ* is a predefined probability density function represented by

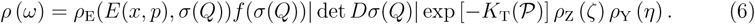

The first two terms *ρ*_E_(*E*(*x, p*), *β*)*f* (*β*) with *β* ≡ *σ*(*Q*) describe the distribution density we want to realize (as explained in Sec. I), which takes the form of a conditional probability. *ρ*_E_(*E*(*x, p*), *β*) describes the density with respect to the PS parametrized by *β*, and the function *f* governs the statistical features of *β*. The variable transformation from *Q* to *β* is needed to construct the ODE using the “unconstrained” variable *Q*, instead of a “constrained” variable such as the inverse temperature *β*, which should be positive. The third term | det *Dσ*(*Q*)| (where *D* denotes the differentiation) in Eq.(6) is the Jacobian needed for this variable transformation to represent integration formulas giving space averages of observables [25, 31], as will be seen in Eq. (15). The remaining terms define the distribution of (*ζ, 𝒫, η*).

The functions *T* (*x, p, Q*), *V*_*E*(*x,p*)_(*Q*), *τ*_Z_ (*ζ*) and *τ*_Y_ (*η*) in Eq. (4) are defined as follows:

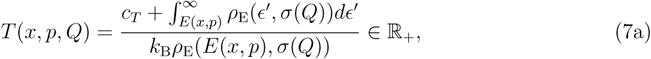

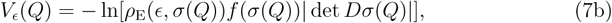

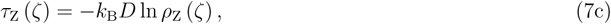

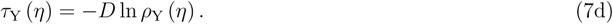

See Ref. [26] for detailed mathematical conditions for these functions. Here, we focus our attention on the *m* = 1 case. For *ρ*_E_, the Boltzmann-Gibbs density for the PS

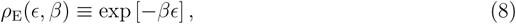

is the main target of our study (but not restricted to it). In this case, *c*_*T*_ = 0 leads to the simple expression

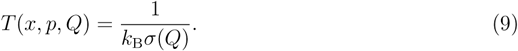

The distribution density function of *ζ, ρ*_Z_ (*ζ*), is typically a Gaussian function, and similar for *ρ*_Y_ (*η*),

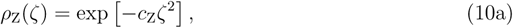

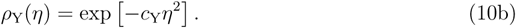

Functions *f* and *σ* are set as follows. *f* : *σ*(ℝ) → ℝ should be strictly positive, of class *C*^2^, and integrable. Usually, a bell type, compact-support distribution is tractable for *f*, and the corresponding *σ* is to be set. Formally, *σ* should be smooth (*C*^3^) and *Dσ* should not be zero, so that the range of *β* becomes an open interval from *β*_L_ ≡ *β*_min_ to *β*_R_ ≡ *β*_max_,

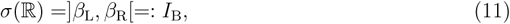

for which we assume 0 ≤ *β*_L_ < *β*_R_ < *∞* for simplicity. Assume that *D* × ℝ^*n*^ × *σ*(ℝ) → ℝ_+_, (*x, p, β*) ↦ *ρ*_E_(*E*(*x, p*), *β*)*f* (*β*), is integrable. For further discussions on *f* and *σ*, see Sections III, IV, and VI A.

### B. Joint distribution for *x, p*, and *β*

From the Liouville equation (5) and the assumption that the vector field *X* is complete, the density *ρ* defined by Eq. (6) becomes the smooth density of an invariant measure with respect to the Lebesgue measure, for the flow generated by *X* [35, 36]. Hence, if the flow is ergodic with respect to measure *P* ≡ *ρdω*, then the long-time average of any *P* -integrable function *g* on phase space Ω gives the space average under density *ρ*, according to Birkhoff’s individual ergodic theorem:

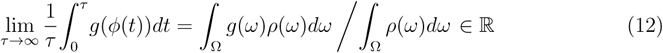

which holds for a solution *φ* of Eq. (4) with an almost every initial point. Thus, for any function of the form *g*(*ω*) = *B*(*x, p, σ*(*Q*)) = *B*(*x, p, β*), we have [26, 31]

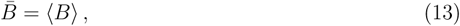

where

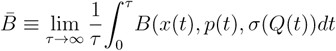

is the time average and

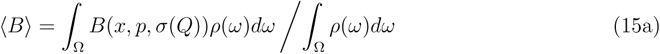

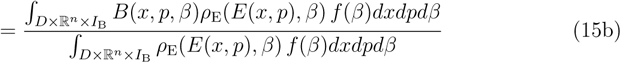

is the space average, assuming that the integrations are finite (such an integrability condition is always assumed hereafter). Namely, (*x, p, β*) is generated with the probability distribution *ρ*_E_(*E*(*x, p*), *β*) *f* (*β*)*dxdpdβ*.

In particular, for a physical quantity *A* : *D* × ℝ^*n*^ → ℝ, we have

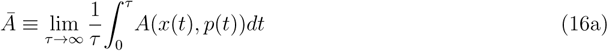

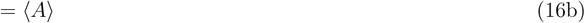

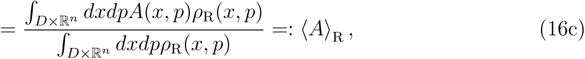

where

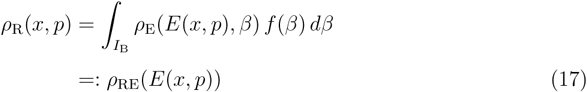

is the (unnormalized) marginal distribution density of (*x, p*). A reweighting to any other density *ρ*_TRG_ : *D* × ℝ^*n*^ → ℝ _+_ can be done by the reweighting formula [26],

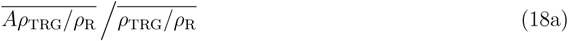

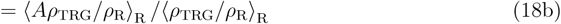

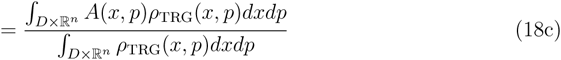

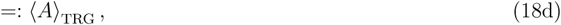

for which *ρ*_TRG_ is typically the BG distribution density *ρ*_TRG_(*x, p*) = exp [−*β*_TRG_*E*(*x, p*)].

On the other hand, for a function of the inverse temperature *h* : *I*_B_ →ℝ, we get

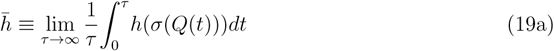

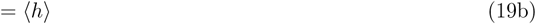

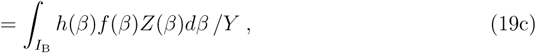

where

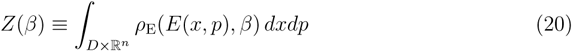

and

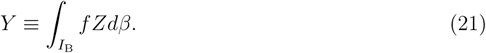

## III. OUTLINE OF THE METHOD

We develop our method introduced in the previous section. The aim is to easily set the desired target distribution of *β* as an input. This is in fact a nontrivial issue since the distribution of *β* becomes

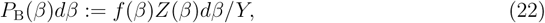

as indicated in Eq. (19) (note that *f* is not equal to the distribution density but the weight), but the partition function *Z* is unknown a priori. If we can set the desired input distribution accurately, we would gain the information of *Z*. It then leads to the accurate evaluation of the space average of *h* represented in Eq. (19), which can be used to check the accuracy of the simulation that yields the time averages of observables only, not the space averages. Furthermore, it significantly helps in the analysis of thermodynamic quantities at every *β* ∈ *I*_B_. However, in a practical sense, it is sufficient to realize the suitable approximation of the desired distribution of *β*, instead of the exact distribution. This is because the expectation of a function of physical variables can be given without any effort via Eq. (16) or (18).

Thus, our specific motivation in the current study is to set in a simple manner the input distribution of *β* that approximately leads to the target distribution. We do this by introducing a suitable input function *f*. Our strategy is simple: estimating an approximation of *Z*, say 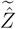, and include this information in the input. Specifically, letting *f*_TGT_ be the desired target distribution of *β*, we define the following *f* :

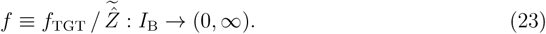

Three general technical remarks are made:

a. The functions *f*_TGT_ and 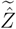 should be strictly positive and smooth (class *C*^2^) to ensure the theoretical condition on *f* (Sec. II A).
b. 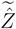 is an approximation of a factorized/normalized partition function:

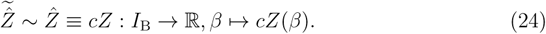 Here, any constant *c* > 0 also becomes a constant factor of function *f* such that it has no effect on the space averages described in Sec. II B, including Eq. (19), via the cancellation. Thus, we can choose the value of *c* in a convenient manner as discussed below.
c. Although *f*_TGT_ can be arbitrarily set in principle, we require that *f*_TGT_(*β*) damps to zero at both ends *β*_L_ and *β*_R_,

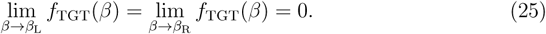

This is useful for the scheme to be robust. Although we may assume 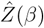 is monotonic with respect to *β*, its specific behavior around both ends *β*_L_ and *β*_R_ is unclear. To ensure the theoretical condition 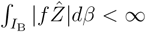 [cf. Eq. (27b)] and conduct a stable simulation with *P*_B_ being finite and having tractable magnitude at both ends, the damping of *f* (*β*) at both ends is needed, which implies the requirement to *f*_TGT_. It would be generally helpful to introduce an additional convergence factor into *f*, especially when the estimation of 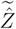 is very rough.

Before discussing the details, we consider the results from Eq. (23). First,

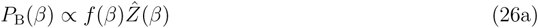

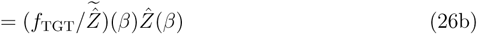

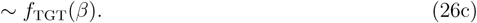

Namely, *P*_B_ is essentially equal to the target distribution *f*_TGT_, and Eq. (19) becomes

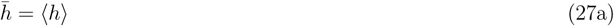

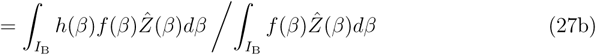

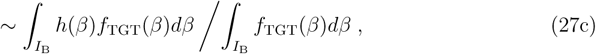

where Eq. (24) is used. This is a very simple formula that can be used in various applications.

The idea of generating the target distribution using a partition function like Eq. (23), is also considered in the simulated tempering (ST) [37, 38]. However, *f*_TGT_ in the current scheme can be freely set under condition (25). The cNH can also provide tempering that is continuous (smooth) with respect to both *β* and the time development. Section IV demonstrates the methods for obtaining the approximation 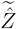 in a simple manner, which is a key technical issue that also has advantages over ST.

Second, *f* constructed as above yields the TS potential (7b) as

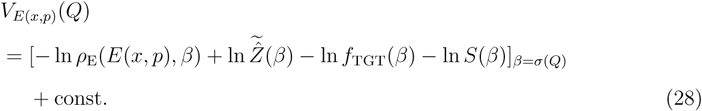

The first term comes from the PS, and describes the interactions between the PS and TS. This term yields just *βE*(*x, p*) if *ρ*_E_ is the case of BG density [Eq. (8)]. The second term comes from the approximation of 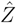 as described above. *f* defined by Eq. (23) is transformed to this term and the third term, which corresponds to a potential function to adjust the shape of the distribution to become the target distribution *f*_TGT_. The final term, defined by

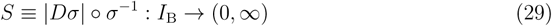

comes from the Jacobian term in Eq. (6), and plays the role of a “repulsion” to keep *β* in the specified region *I*_B_.

The resulting TS potential (28) along with the current cNH formula coupling both the PS and TS yields very original dynamics of continuous variables. It also presents the characteristic features of PS sampling. Section V demonstrates in detail the mutual dynamics between the PS energy *E*(*x, p*) and dynamical inverse temperature *β*.

## IV. DETERMINING THE WEIGHT OF THE *β* DISTRIBUTION

### A. Protocols

To complete *f* defined in Eq. (23), we describe two approaches to construct 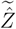. The first approach involves the integration of the energy averages, as detailed in Sec. IV B; the second is a refinement approach detailed in Sec. IV C. This subsection gives brief explanations of these approaches and total protocols by combining the approaches. Although these approaches can be extended to the case of any *ρ*_E_ [see e.g., Eq. (30) below], we here describe the BG case [Eq. (8)] for simplicity and use terminologies such as “energy average,” instead of “energy-like quantity average”, which can be used in extended cases. Specifically, the energy average here is the phase-space average (expected value) of the PS total energy under the BG distributions, ⟨*E*⟩_BG;*β*_.

The first approach constructs 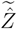 by taking in the energy average data 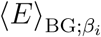 at selected individual temperatures 1*/k*_B_*β*_*i*_ for *i* = 1, 2, …. These data can be obtained by simultaneously performing BG MDs, such as the NH EOM. An integration formula with respect to these data is derived. In this procedure, a smooth function 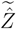 [see Sec. III, (a)] can be obtained by an interpolation method.

However, it should be noted that the energy averages obtained with this approach may not be accurate owing to insufficient sampling with the BG MDs. Thus, we take the second approach, where 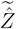 is refined by a simple correction. The proposed correction is to eliminate the deviation of the output distribution from the input.

Therefore, the proposed simple protocol to construct 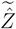 is as follows:

I. Perform a BG MD at temperature 1*/k*_B_*β*_*i*_ and calculate 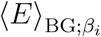 for each *i*.
II. Define 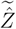 by using 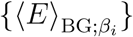.
III. Perform the cNH.
IV. Redefine 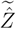 by a correction. Some optional variations are discussed here. First, in (I), the energy averages may be estimated by applying a single enhanced sampling method, such as the cNH, along with the reweighting formula (18). However, a certain knowledge of the target system should be obtained in advance. Second, instead of (IV), one can repeat the first approach, i.e., calculating the BG energy averages via the reweighting from the output of the cNH in (III), and then defining 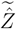 using these energy averages. The 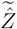 obtained via enhanced sampling would be a better approximation than that obtained in (I) and (II). Although the first approach is universal and can be used as both an initial guess and refinement to construct 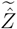, the second approach in (IV) is recommended for the refinement owing to its simplicity, as detailed in Sec. IV C. After the protocols (I)–(IV) for calculating 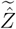,
V. perform the cNH again,

as the *production* run. In summary, the current protocol includes numerical simulations of (I) the BG MD runs, (III) the cNH *preliminary* run using the approximated 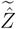 in (II), and (V) the cNH production run using the refined 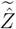 obtained in (IV).

### B. Construction of 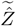 using energy averages

We begin by observing the relation between *Z* and the average of an energy, or generally an energy-like quantity [see Eq. (31)]. For a general *ρ*_E_, we see

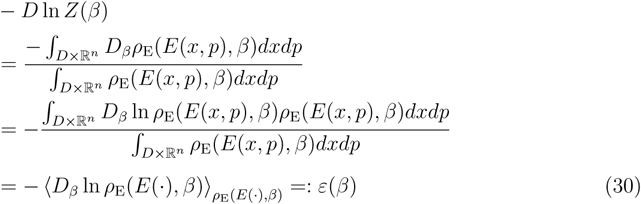

for each *β* ∈ *I*_B_, noting that

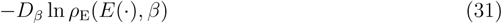

can be seen as the energy-like quantity since it has a dimension of the energy (*D*_*β*_ denotes the partial differentiation with respect to *β*). In fact, in case where *ρ*_E_ is the BG density (8), it follows that

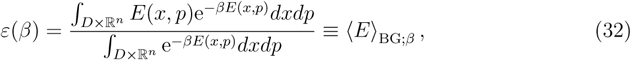

viz., *ε*(*β*) becomes the (total-)energy average under the BG distribution at temperature 1*/k*_B_*β* for every *β*. Integration of Eq. (30) with respect to *β* leads to

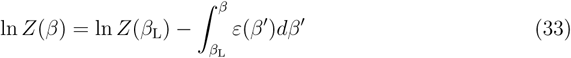

or

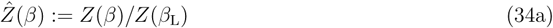

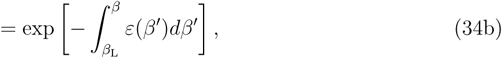

which corresponds to the fact that the constant in Eq. (24) is chosen as *c* ≡ 1*/Z*(*β*_L_). Equation (34), named herein as the “integration formula”, is the exact relationship between the partition function and the average of the energy (or the energy-like quantity, in general). The relationship (33) was approximated in previous approaches (Taylor expansion [37] and a modified linear relation [39]) using discretized *β*. The exact relationship (33) or (34) in the current scheme significantly helps our treatment and enables us to obtain an good initial guess for 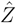.

#### Remark 1

Note again that 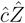 suffices for 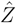 for any *č* > 0, since *ĉ* becomes a constant factor of f.

The approximation of 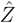 proposed in Protocol (II) is based on the assumption that we have data *e*_*i*_ that approximate values *ε*(*β*_*i*_) (*i* = 1, …), and is based on spline-function interpolation using these data. Given {*β*_*i*_, *e*_*i*_}_*i*=0,1*…,M,M*+1_, where *β*_*i*_ ∈ *I*_B_ =]*β*_L_, *β*_R_[is discrete data of the inverse temperature such that 0 *< β*_L_ ≡: *β*_0_ *< β*_1_ *< β*_2_ *<…< β*_*M*_ *< β*_*M*+1_ :≡ *β*_R_, and *e*_*i*_ approximates *ε*(*β*_*i*_) as

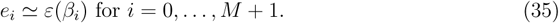

For convenience, we define bins

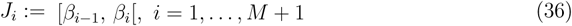

and *h*_*i*_ ≡ |*J*_*i*_| = *β*_*i*_ − *β*_*i*−1_ for *i* ∈ ℳ ≡ {1, …, *M* + 1}. By using only these data {*β*_*i*_, *e*_*i*_}, along with additional conditions (see Appendix A), we define a spline function

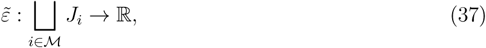

and an approximation of 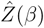:

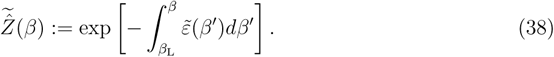

We define 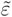 as piecewise on each *J*_*i*_ by

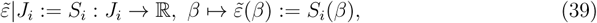

where *S*_*i*_ (*i* ∈ ℳ) is a polynomial with a specific form according to the scheme of interpolation. In Appendix A, we present the cubic (third-order) spline interpolation scheme, and summarize the resulting specific forms of quantities employed in the current method. The cubic scheme, yielding *S*_*i*_ as a third-order polynomial, is simple and has fine properties, although the current method is not limited to a specific kind of interpolation.

### C. Correction of 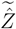

If the phase-space sampling in the cNH preliminary run [Protocol (III)] is sufficient, then the output distribution density of *β, P*_B:out_(*β*), gives a good approximation of *P*_B_(*β*) [see Eq. (26a)] such that

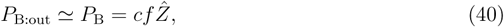

where *c* is the normalization constant. It should be noted that the error of this approximation mainly comes from the discrepancy between 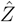 (which is exact but unknown) and 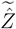 (which is used as the input of the preliminary run), but not from the finite-time sampling (under the ergodicity assumption). Thus,

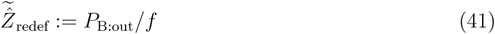

gives a better approximation of 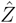 than that previously defined. The constant *c* can be ignored for defining 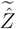, as stated in Remark 1. We note that if the sampling is complete, then this 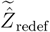 gives the exact 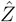 (but also see below). This is an idea of Protocol (IV) demonstrated in Sec. III. A technical issue arises in which *P*_B:out_ is obtained as numerical discrete data, and not as a continuous function. This can be resolved by employing the spline-function interpolation, as described above.

If we want to have *P*_B:out_ that is very similar to *P*_B_, we can iterate the process of this kind of redefinition of 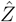 That is, the (*k* + 1)-th approximation 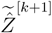 can be defined by induction using the *k*-th approximation 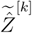 and distribution output 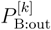 in the *k*th run:

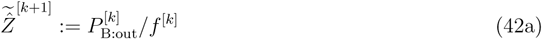

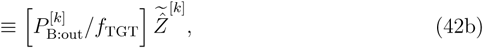

where Eq. (41) is read as the initial process *k* = 1 with 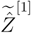. We see that

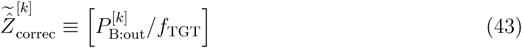

is the correction factor from the *k*-th to the (*k* + 1)-th approximation: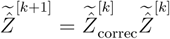.

Again, it should be stressed that this iteration is not necessary when the approximate distribution 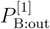 is sufficient in practice. In fact, the average of any physical-space quantity *A*(*x, p*) in the target distribution is obtained by the reweighting formula [Eq. (18)], so that it can be accurate despite 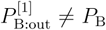. What we have to do is to realize a *β*’s distribution that is tolerably similar to the target distribution *f*_TGT_ in a major region of *β*, instead of specifying the exact *β*’s distribution proportional to *f Z*.

Although the cNH differs from the McMD in many respects, regarding the potentially iterative procedure (42) only, its mathematical structure is essentially the same as that taken in the McMD method [10]. The difference is in the iteration target, which is the energy density of state Ω(*ϵ*) in the McMD, instead of 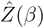. Note that 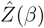 obtained in the current scheme will be useful to obtain information on Ω(*ϵ*) through the Laplace transform. Finally, it should be noted that the correction procedure proposed here can be applied to any *ρ*_E_, and is not limited to the BG case.

## V. RESULTING TS POTENTIAL AND SAMPLING PS STATES

We describe the TS potential and TS force, which are specific to the cNH and characterize the sampling features. We consider here the typical case in which *ρ*_E_ is the BG; a general case is discussed in Appendix B.

The TS potential can be written, up to constant, as

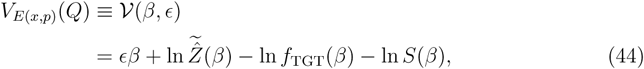

where *ϵ* = *E*(*x, p*) and *β* = *σ*(*Q*). The graph for each potential term is schematically shown in Fig. 2. The corresponding TS force becomes

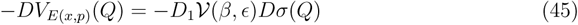

with

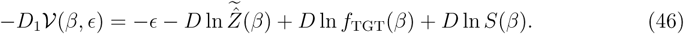

**FIG. 2.**
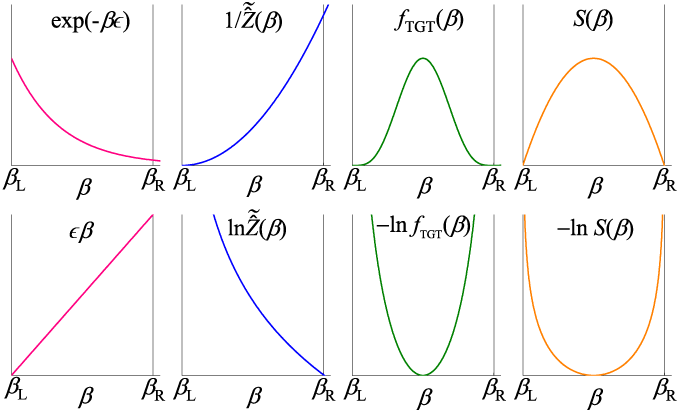
Schematic figure of each term of the TS potential [Eq. (44)] (bottom panel) and its original quantity (top).

Note that although the TS potential and force actually act on *Q*, instead of *β*, in the EOM, the fact that *Q* and *β* correspond one-to-one enables us to identify *β* with *Q*. Thus, we here consider the motion of *β* in its space *I*_B_, since this viewpoint helps us catch the essential features of cNH dynamics. In line with this, potential (44) in the TS should be described by *β*, while *ϵ* is a parameter. However, *ϵ* = *E*(*x, p*) is not a static parameter because its value changes during the simulation. Thus, it is useful to consider the TS potential in the 2D (*β, ϵ*)-space, such as *𝒱*(*β, ϵ*). It should also be noted that, since *E* is a parameter for the TS, the TS does not feel the force along the *ϵ*-direction; it only feels the force along the *β*-direction, as described by Eq. (46). Thus, the stationary point of the TS is determined by the (partial) derivative of the potential with respect to *β*. To clarify the discussions, we assume the following:

### Assumption 1.

⟨*E* ⟩_BG;*β*_ *is strictly monotonic decrease with respect to β*.

### A. Major terms of the TS potential: Shape

We begin by considering only the first two terms of the potential, i.e.,

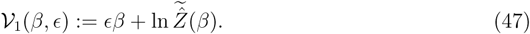

Then, the stationary point of *𝒱*_1_ is characterized by

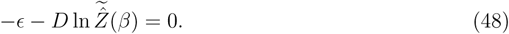

This stationary point *β* moves according to the movement of *ϵ* = *E*(*x, p*) (see Fig. 3). Thus the set of the stationary points

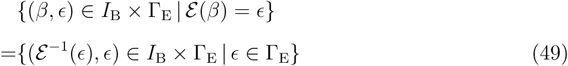

**FIG. 3.**
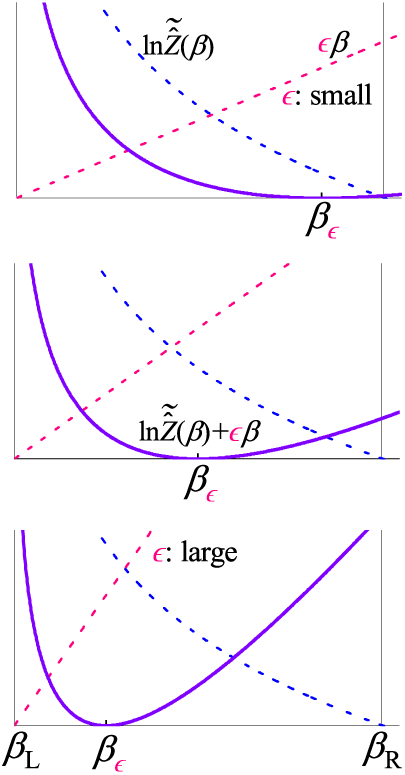
Schematic figure of the two major terms of the TS potential and their summation [with the shift via Eq. (61)]. The stationary point *β*_*ϵ*_ moves according the increase in *ϵ* (from top to bottom panel).

is defined. Here, two assumptions have been made that the function

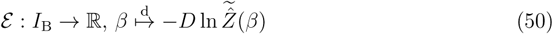

is injective and that an effective energy region observed in the simulation, Γ_E_, meets the condition,

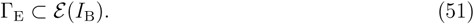

It is natural to define Γ_E_ as a region characterized by two energy averages such as

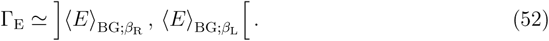

These two assumptions ensure that the stationary point uniquely exists for each energy value in Γ_E_. They are physically reasonable by considering Eqs. (30) and (32), since we have

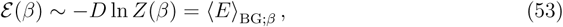

indicating that *ε* ∼⟨*E* ⟩_BG;(·)_ is injective owing to Assumption 1, and since we thus have

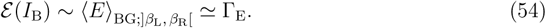

The reason why definition (52) is natural is based on the fact that the most probable, or the representative, energy value at a given *β* value is near the energy average under the BG distribution:

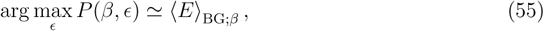

where *P* (*β, ϵ*) is the probability density on the 2D (*β, ϵ*)-space. To see Eq. (55), first note that *P* (*β, ϵ*) becomes proportional to

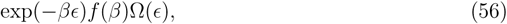

where Ω(*ϵ*) is the density of state for the total energy. Thus, the probability density for which the total energy attains *E*(*x,p*) = *ϵ* under the condition that *β* =constant is proportional to

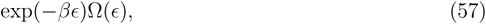

which is equal to the probability density for *E*(*x, p*) = *ϵ* under the BG distribution at temperature 1*/k*_B_*β*. For every *β*, thus arg max_*ϵ*_ *P*(*β, ϵ*) = arg max_*ϵ*_ exp(−*βϵ*)Ω(*ϵ*), which implies Eq. (55) in normal PSs.

Taking *β* as an independent variable, the set (49) can be represented as

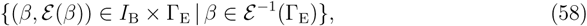

which is now a curve parameterized by *β*, and then approximated by the curve

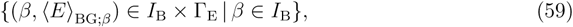

as introduced in Sec. I [note that a set {(*β*, ***ε*** (*β*)) *∈*ℝ^2^ | *β ∈ I*_B_}, which is a minor extension of set (58), makes sense even if Γ_E_ ⫋ ****ε**** (*I*_B_)]. The trajectory of the TS should lie along such a curve, which seems like a “river” flowing through the “valley” *𝒱*_1_ [see Fig. 4(a)]. The reason why *𝒱*_1_ makes a valley comes from the fact that the stationary point is in *I*_B_ [as in Eq. (58)] and that *𝒱*_1_ should be concave with respect to *β* for any *ϵ ∈* Γ_E_:

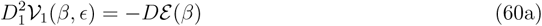

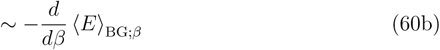

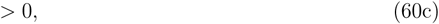

where Assumption 1 is again used in the last line.

**FIG. 4.**
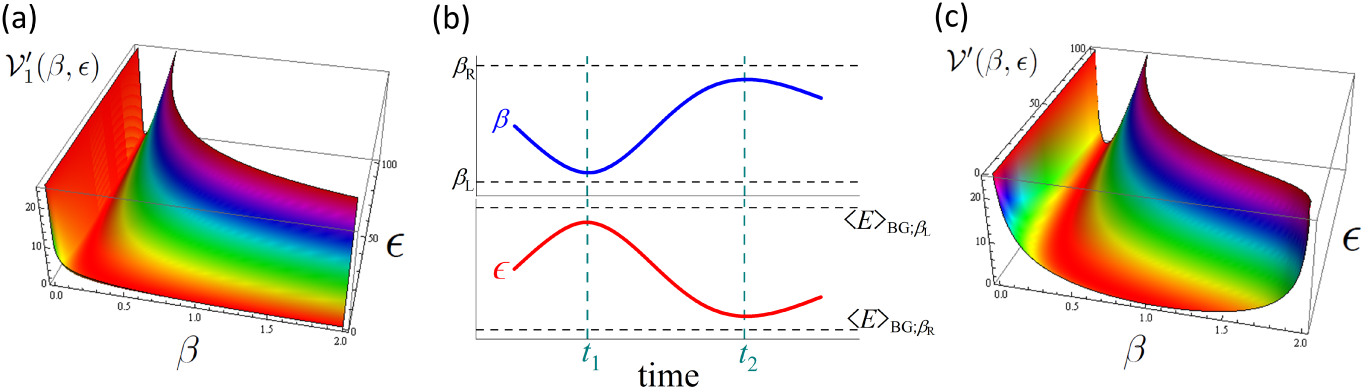
(a) TS potential described by the first two terms. Indicated are that for the 2HO model system described in Sec. VI. (b) Schematic graph indicating the processes (i) and (ii) regarding the dynamics of *β* and *ϵ* = *E*(*x, p*). The behavior before *t* = *t*_1_ indicates the process (i) *E* grows ⇒ *β* decays ⇒ *ϵ* grows. At *t* = *t*_1_ it switches into process (ii), where *ϵ* decays ⇒ *β* grows ⇒ *ϵ* decays. At *t* = *t*_2_, it again switches into (i). (c) TS potential described by all the terms for the model system in (a). The shift defined by Eq. (72) is used to depict these TS potentials.

Since the gradient with respect to the *ϵ*-direction can be ignored in the TS, we can use, instead of *𝒱*_1_, a shifted potential defined by

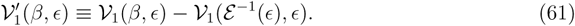

Here, the shifting, which is irrelevant to *β*, does not affect the TS force. In addition, the stationary point *β*_*ϵ*_ ≡ ***ε***^−1^(*ϵ*) remains (since 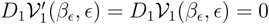, where

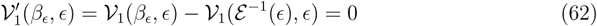

holds, viz., the “river” lies at the height of zero, which helps us understand the features of the potential.

### B. Sampling dynamics

The movement of *ϵ* = *E*(*x, p*) shifts the stationary point for *β* and so induces the movement of *β* along the river. As seen above, *β* is stable if *E*(*x, p*) is around its average ⟨*E*⟩_BG;*β*_. However, small oscillations of *β* around its certain values in the river do not result in enhanced sampling. The cNH provides two sampling mechanisms. To see the first one, we begin with the fact that an increase in *ϵ* induces the decrease in *β*, and vice versa. These directions of motion are derived from the two terms of the TS force (46):

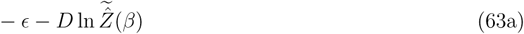

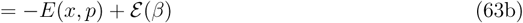

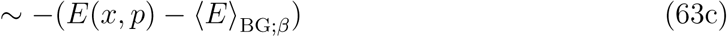

[note that this increase-decrease relationship between *E*(*x, p*) and *β* is irrelevant to the signature of *Dσ*, which translates the TS force acting on *β* into that on *Q*, as seen in Eq. (45)]. On the other hand, the movement of *β* induces the movement of *ϵ* = *E*(*x, p*). This is because, if *β* decays [grows], which indicates the increase [decrease] in the dynamical temperature *T*_D_ ≡ *T* (*x, p, Q*) = 1*/k*_B_*σ*(*Q*) = 1*/k*_B_*β* [see Eq. (9)], then the PS energy *E* grows [decays] in most cases. Hence, the cNH provides the first mechanism such that

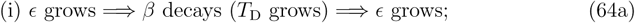

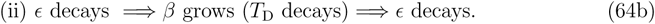

Each process indicates only the monotonic change: *ϵ* simply grows and *β* simply decays in (i), which becomes an “accelerate” process considering the fact that *β* decay indicates the *T*_D_ growth; and vice versa in (ii), which becomes a “decelerate” process. This *mutual boost* between *ϵ* and *β* is a remarkable feature of the sampling in the cNH, in contrast to the conventional feedback mechanism around a single equilibrium point.

However, each process does not continue infinitely. The second sampling mechanism is a switching due to an allowance of perturbation via the fluctuations. In fact, the *β* decay in (i) induces not only the *ϵ* growth, but also *ϵ* decay occasionally. In the latter case, process switches to process (ii). Conversely, switching from (ii) to (i) is also possible.

Namely, the “accelerate” process (i) or “decelerate” process (ii) continues for a relatively long time, and they occasionally switch. Such a move is schematically shown in Fig. 4(b). This is a fundamental mechanism of (*β, ϵ*)-space sampling in the cNH, where *T* -space sampling and the *E*-space sampling mutually cooperate. It also provides a specific mechanism that explains the features of enhanced sampling, as often intuitively stated in generalized sampling methods, such that the *T* -space “random walk” enhances the *E*-space “random walk.” Note that the movement of *β* from *β*_L_ to *β*_R_ corresponds to the movement of the energy from 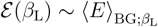 to 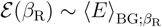.

### C. All terms of the TS potential

All the potential terms of *𝒱* defined in Eq. (44) should be taken into account. Basically, this implies the deformation or perturbation of the shapes of the valley and river discussed so far, which will become

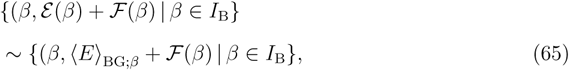

where ℱ (*β*) is the sum of the last two terms of Eq. (46). To understand this implication, we specifically add the logarithmically-concave condition of *f*_TGT_*S*:

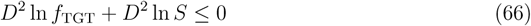

(obviously, the logarithmically-concave conditions of individual two functions suffice). Then

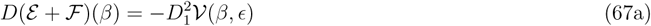

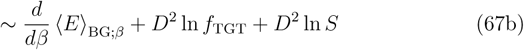

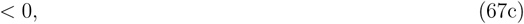

indicating that *ε* + ℱ is also injective, so that the discussion in section V A still holds. The total TS potential *𝒱* is also indicated to be convex with respect to *β* and makes a valley.

The third term, − ln *f*_TGT_(*β*), of *𝒱* plays the role of making adjustments to build the desired distribution of *β*. It is compatible with the purpose of realizing any distribution and the requirement (25) on the zero damping at the ends. This is because any non damping density 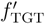 on any 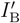 can be smoothly extended to a damped function *f*_TGT_ defined on 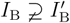 (i.e., 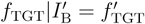) in many cases, indicating that *f*_TGT_ can be viewed as a new target.

The last term − ln *S*, which originated from the Jacobian term, seems to be a “restraint” potential of the TS. To see this explicitly, we add a condition for *σ* such that

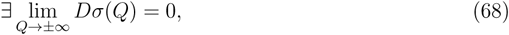

which is natural (but not ensured by the other conditions) in the sense that the region of *β* is assumed to be bounded. Then, we can show that *S*(*β*) damps to zero at the both ends of *β*, and that the last term becomes a “restraint” potential, i.e.,

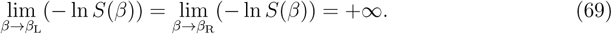

The last term keeps the “U”-like shape (see Fig. 1) when

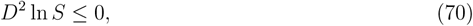

or, equivalently, *DσD*^3^*σ ≤* 2(*D*^2^*σ*)^2^. See Fig. 4(c) which describes the total potential *𝒱*(*β, ϵ*).

In addition to the last term, the third term also becomes a restraint owing to the damping of *f*_TGT_ at both ends. These restraints are useful because the movement derived from (i) and (ii) in Eq. (64) should terminate at the ends *β*_L_ and *β*_R_. Namely, when *β* arrives near *β*_R_, the repulsive force acts through these restraint potentials, and the trajectory returns, implying that process (ii) switches to process (i). A similar mechanism with the opposite direction holds near *β*_L_.

First, it should be noted that, as a counterpart of Eq. (51), we must show

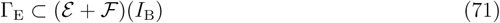

to keep the discussion in section V A. Equation (71) is valid since (*ε* + ℱ)(*I*_B_) = ℝ, which can be derived from the fact that Eq. (69) leads to *D* ln *S*(*I*_B_) = ℝ. Second, the shifting method in Eq. (61) can also be utilized for each term of the TS potential and for all the terms such as

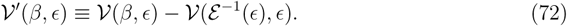

Finally, we note that the addition of ℱ (*β*) in the TS-force [Eq. (63)] plays the role of perturbation in the process “*E* grows [decays] ⇒ *β* decays [grows]” in Eq. (64).

## VI. NUMERICAL SIMULATION

We applied the current method to two systems: a low dimensional model system and a realistic bimolecular system. In Sec. VI A, after demonstrating common simulation conditions between the two systems, we describe the motivations and simulation details for individual systems. The simulation results are shown in Sec. VI B.

### A. Simulation condition

We set *ρ*_E_ to be the BG density (8) and set *ρ*_Z_ and *ρ*_Y_ to be Gaussian functions (10). The function forms of *σ* and *f*_TGT_ are defined as follows. A sigmoid function was used for *σ* defined as

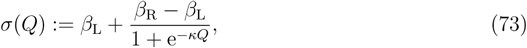

since it is easy to set the range of *β*, which becomes]*β*_L_, *β*_R_[= *I*_B_. Here, *κ* > 0 is an arbitrary nondimensional parameter, and the other constants are set in such a way that *Q* = 0 corresponds to the average inverse temperature,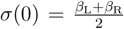. All the conditions required for *σ* are met, and the logarithmically-concave condition (70) is valid, where

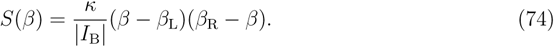

On the basis of the Beta distribution, we use

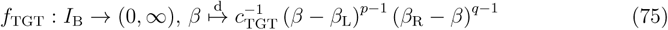

for *p, q* > 1, where 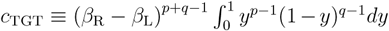 is the normalization constant. Clearly, *f*_TGT_ damps at both ends, and the logarithmic concavity *D*^2^ ln *f*_TGT_ ≤ 0 also holds. Thus, Eq. (66) is valid. We set the symmetric condition, *p* = *q* = 5.

To numerically integrate the ODE (4), an explicit, symmetric, second-order integrator was used, as described in Appendix C. Based on the extended space formalism [40, 41], the invariant function [see Eq. (108)] enables us to easily check numerical integration errors.

#### 1. 2HO system

As a first example, we utilized an *n* = 2 dimensional harmonic oscillator (2HO) system defined by the energy function

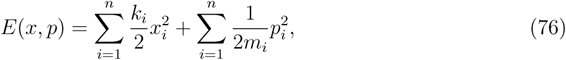

where *k*_*i*_ and *m*_*i*_ are the spring constant and mass for the *i*-th degree-of-freedom, respectively. All quantities were treated as dimensionless.

##### Motivations

(i) This model is fundamental because it is the simplest model that describes PS behavior around a stable state. (ii) It is suitable for validating the current method, since the exact partition function and (marginal) distributions are known in this system. The exact partition function *Z*(*β*) ∝ *β*^−*n*^ leads to the setting

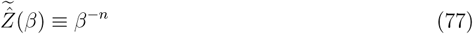

and enables us to rigorously check the distribution of *β* generated by the current method, where the exact distribution density is

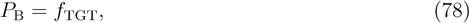

following from Eqs. (22) and (77), and from the fact that both *P*_B_ and *f*_TGT_ are normalized. Distribution densities for *P, ζ*, and *η* are clear, while that for *ω*_*i*_ = *x*_*i*_ (*i* ≤ *n*) or *p*_*i*−*n*_ (*i* > *n*) with the use of Eqs. (8) and (75) is given by

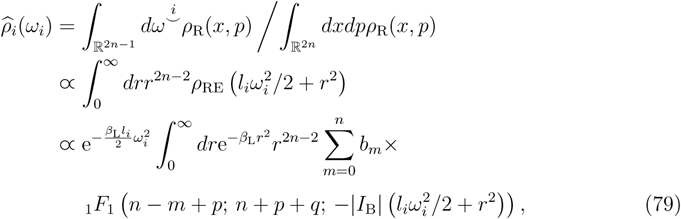

where 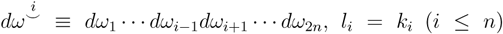 or 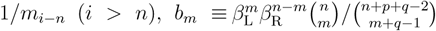, and _1_*F*_1_ is the confluent hypergeometric function. We get 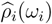 by normalizing Eq. (79) and numerically estimating 1D integrations. Assumption 1 (Sec. V) is valid because ⟨*E*⟩_BG;*β*_ = *n/β*. (iii) It is interesting to see the ergodicity in the cNH and obtain the accurate target distribution, since some sampling methods, e.g., the conventional NH EOM, fail for *n*HO [42, 43]. (iv) It becomes normal by taking *n* ≥ 2 in that the density of state Ω(*ϵ*) = *cϵ*^*n*−1^ (*c* is irrelevant to *ϵ*) increases with respect to the total energy *ϵ*. In contrast, the 1HO, in the case of *n* = 1, gives Ω(*ϵ*) = constant and does not allow compatibility with assumption (55), as arg max_*ϵ*_ *P* (*ϵ, β*) = arg max_*ϵ*_ exp(−*βϵ*) = 0 for any *β*. Thus, we used the minimum compatible value *n* = 2.

##### Conditions

We used the following parameter values: *k*_B_ = 1, *k*_1_ = *k*_2_ = 1, *m*_1_ = 1, *m*_2_ = 2, *β*_L_ = 0, *β*_R_ = 2, *κ* = 1 [Eq. (73)], *c*_Z_ = *c*_Y_ = 1 [Eq. (10)], and **M**_T_ = 1 [Eq. (4)]. The initial values *x*_1_(0) = *x*_2_(0) = 0, *p*_1_(0) = 1, *p*_2_(0) = −1, *ζ*(0) = 0, *Q*(0) = 0, *𝒫*(0) = 1, and *η*(0) = 0 were used for 10^8^ timestep numerical integration using the unit time of 1 × 10^−3^.

#### 2. Chignolin system

The second example is chignolin in explicit water with the all atom descriptions. Chignolin is a ten-residue designed peptide that forms a stable (beta)-hairpin structure [44].

##### Motivations

(i) This system is typically used for the simulation study of proteins; hence, the system can be treated as a benchmark test. In fact, it has been frequently used to investigate various molecular simulation methods [45–49]. (ii) In contrast to the *n*HO system, 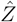 is unknown in this realistic system. Thus, we can check the validity of all our existing schemes, viz., the construction method of 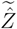 for the preliminary run and its correction method for the production run. (iii) We show that the desired wide inverse-temperature distribution can be easily realized with sufficient accuracy for this typical system. (iv) We clarify two algorithmic advantages. In contrast to many sampling methods, the convergence of the simulation data can be checked by simply monitoring the distributions of *ζ, 𝒫*, and *η*, since their exact distributions are known. In contrast to many stochastic methods, the accuracy of the numerical integration of the fundamental EOM can also be checked by monitoring the value of the invariant function (108). Note that good results from these two kind indicators will be the necessary conditions for sampling convergence and integration accuracy, respectively, but will not be the sufficient conditions (which are not obtained in general cases). (v) We investigate the sampling efficiency in the PS by calculating free energy landscapes, and examine the unique and interesting sampling mechanisms discussed in Sec. V.

##### Conditions

The initial structure of the chignolin molecule was taken from NMR structures in the PDB entry 1UAO. The solution system consists of 1941 TIP3 water molecules [50] with two sodium ions for electrically neutralizing the system. The chignolin molecule and solution system (total 5963 atoms) were contained in a rectangular MD cell, and were equilibrated under the *NTP* ensembles in advance. Numerical simulations for cNH EOM (4) were performed with the timestep of 1 fs using the AMBER ff03 force field [51]. After the preliminary run [Protocol (III)] of the cNH 1-*µ*s simulation, the production run for 1-*µ*s was conducted [Protocol (V)], as described in Sec. IV A. For the initial values of these two runs, the equilibrated values described above were used for *x*(0) and *p*(0), while *ζ*(0) = *Q*(0) = *P*(0) = *η*(0) = 0. We set the range of *β* so that it covers the temperature from 240 K to 600 K, via 1*/k*_B_*β*_L_ = 600 K (*β*_L_ = 0.839 [kcal/mol]^−1^) and 1*/k*_B_*β*_R_ = 240 K (*β*_R_ = 2.097 [kcal/mol]^−1^). The reweighting to the BG distribution density was done by Eq. (18) via *ρ*_TRG_(*x, p*) ≡ exp[−*β*_TRG_*E*(*x, p*)]. The parameters used were *c*_Z_ = 10^−3^ [ps kcal/mol]^−2^ [Eq. (10a)], *c*_Y_ = 10 ps^−2^ [Eq. (10b)], **M**_T_ = 3 × 10^−3^ ps^2^ [Eq. (4)], and *κ* = 10^−2^ [Eq. (73)].

### B. Numerical Results and Discussion

#### 1. 2HO system

Figure 5 shows the distribution densities generated by the cNH with respect to coordinate *x*, momentum *p*, and control variable *ζ* for the PS. The simulated results and (exact) theoretical results showed good agreement. The standard deviations of the discrepancies between the simulated and theoretical densities were 2.1 × 10^−3^, 1.9 × 10^−3^, 2.1 × 10^−3^, 1.4 × 10^−3^, and 1.1 × 10^−3^ for *x*_1_, *x*_2_, *p*_1_, *p*_2_, and *ζ*, respectively. This implies that the ergodicity of the total system was attained, which is a nontrivial task for sampling methods, as shown in Fig. S1. It should be a stronger property than that required for “enhanced” sampling methods. As expected, the trajectories of *x* and *p* exhibit ergodic features, as shown in Figs. 6 and S2. As the duration of trajectories increased, large fluctuations as rare events increased and their amplitudes seem to be enlarged. In contrast, the invariant function remained constant, indicating that the numerical integration of this chaotic system was performed accurately.

**FIG. 5.**
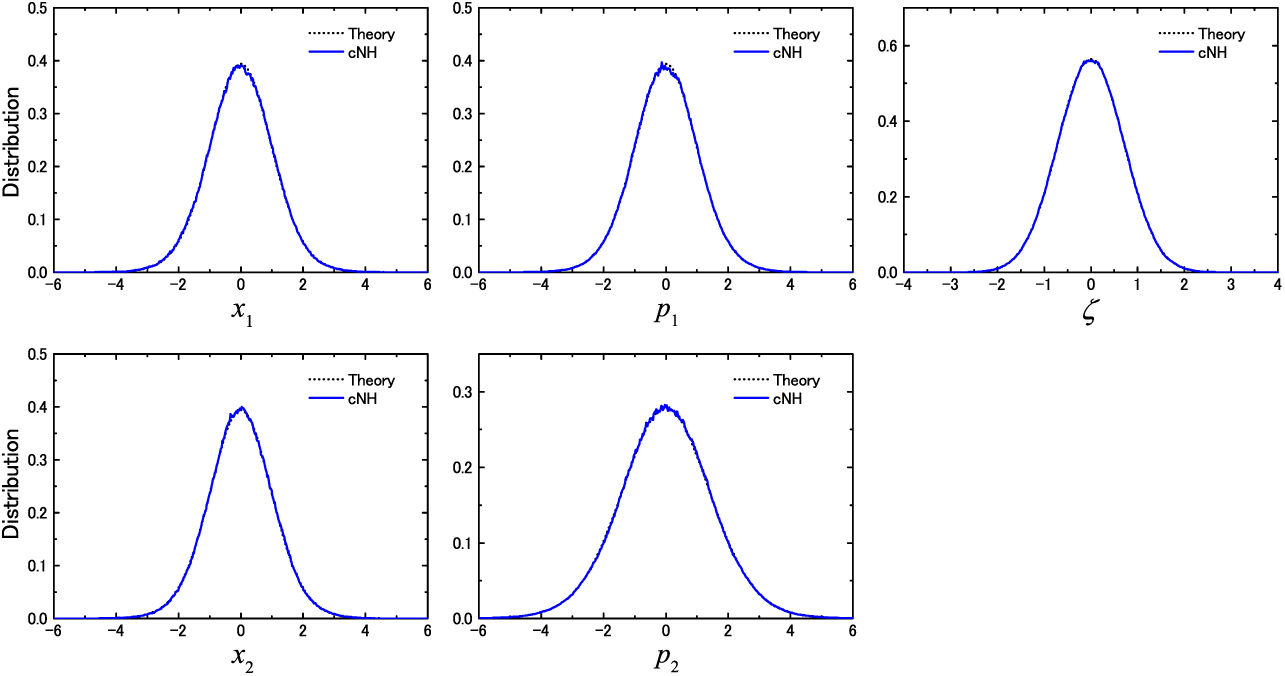
1D marginal distribution densities of *x, p*, and *ζ* for the 2HO model. The theoretical density and simulated density obtained by cNH are shown by broken and solid lines, respectively.

**FIG. 6.**
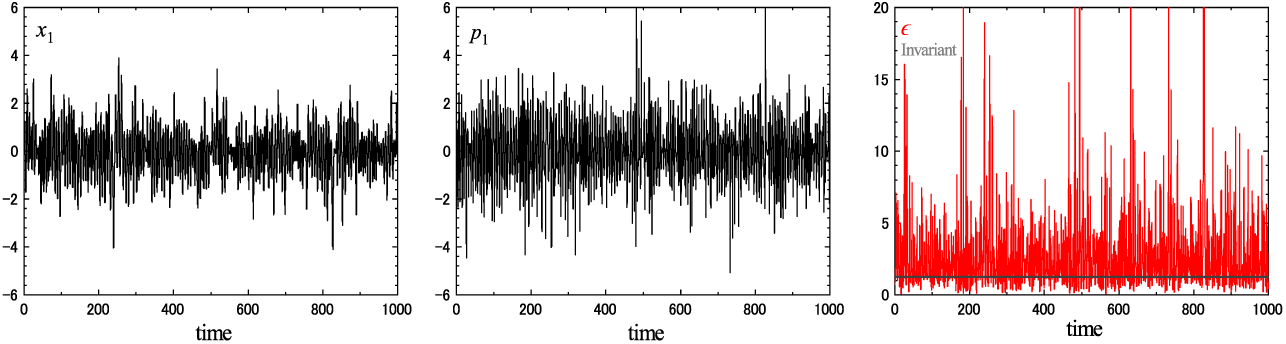
Time courses of *x*_1_, *p*_1_, and *ϵ* = *E*(*x, p*), and the invariant for the 2HO model. Behaviors in various time durations are shown in Fig. S2.

Figure 7(a) shows the distribution density with respect to *β*, viz., *P*_B_. This should be equal to *f*_TGT_ theoretically [Eq. (78)], as confirmed in the figure. In addition, we compared the simulated and theoretical distribution densities of the remaining TS variables, viz., momentum *𝒫* and control variable *η* [Figs. 7 (b) and (c)]. The standard deviations of the discrepancies were 6.1 × 10^−3^, 1.1 × 10^−3^, and 1.2 × 10^−3^ for *β, 𝒫*, and *η*, respectively. These discrepancies were smaller for longer simulations. Hence, we conclude that the ergodic condition is valid, the simulation results were accurate for the 2HO system, and the computational scheme (23) is appropriate.

**FIG. 7.**
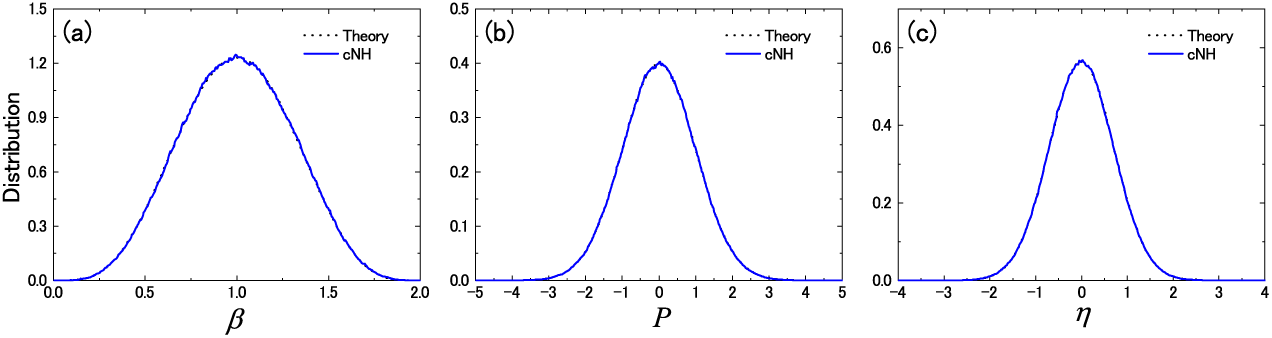
Distribution densities of (a) *β*, (b) *𝒫*, and (c) *η* for the 2HO model. The theoretical and simulated densities are shown by broken and solid lines, respectively.

These enhanced samplings were realized from the mutual dynamics of the dynamical inverse temperature *β* and PS energy *ϵ* = *E*(*x, p*), as characterized by processes (i) and (ii) along the minimum path (viz., the “river”) made by the TS potential, as discussed in Sec. V. In fact, the “schematic” graphs for the TS potential in Figs. 2, 3, 4(a), and 4(c) are the real curves (using function rescaling and the shifting) for this 2HO system. Figure 8(a) shows the minimum path of the TS potential and the (*β, ε*) plots generated by the simulation, indicating that the dynamics are explored around the path, where the reduced attainment for *β* ∼ 0 is due to the restraint term of the TS potential. Figures 8(b) and 8(c) show the individual trajectories of *ε* and *β*. First, we admit the behavior, in a certain period, characterizing the mechanism such that *E* grows [decays] ⇔ *β* decays [grows]. For example, see Fig. 8(c) for the characteristic behavior in the accelerate process (i) observed for a certain duration before *t* = *t*_1_, where *ε* grows and *β* decays. Then, it switches into decelerate process at *t* = *t*_1_ and continues the process up to *t* = *t*_2_. Afterward, it again switches into (i). The perturbation via the fluctuations of *E* and the effect of the additional terms in the TS potential cause these switches and generate nonlinearly oscillating trajectories.

**FIG. 8.**
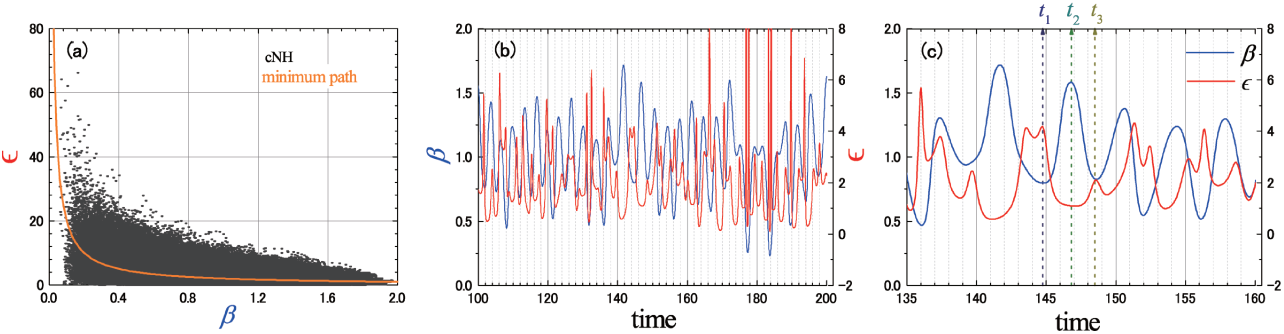
(a) The minimum path (or “river”) made by the TS potential on the 2D (*β, ϵ*)-plane and the scatter plots generated by cNH for the 2HO system. (b) Time courses of *β* and *ϵ* = *E*(*x, p*) for cNH of the 2HO. (c) Close-up view of (b): the accelerate process (i) is observed for a certain duration before *t* = *t*_1_, where *ϵ* grows and *β* decays (*T*_D_ grows); it switches at *t* = *t*_1_ into the decelerate process (ii), where *ϵ* decays and *β* grows (*T*_D_ decays); at *t* = *t*_2_ it again switches into the accelerate process (i) and continues up to *t* = *t*_3_.

#### 2. Chignolin system

We follow the scheme outlined by Protocols (I)–(V) stated in Sec. III. As Protocol (I) for the preparation to construct 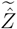, we set seven *β*_*i*_ at almost even intervals from *β*_L_ to *β*_R_. That is, *M* = 5 internal points in Eqs. (35) and (36), where *ε*(*β*_*i*_) = ⟨*E*⟩_BG;*βi*_ is the energy average under the BG distribution at temperature 1*/k*_B_*β*_*i*_, and its approximation *e*_*i*_ was evaluated by a preceding MD simulation using the NH EOM for each *i*. The data imply that Assumption 1 holds in this system. Then, as Protocol (II), according to the spline function scheme described in Sec. IV B, we obtained a smooth function 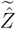 [see Eq. (38) and Eq. (86)] on *I*_B_. These results are shown in Fig. 9. Hence, we had all necessary input data to conduct the cNH simulation in Protocol (III).

**FIG. 9.**
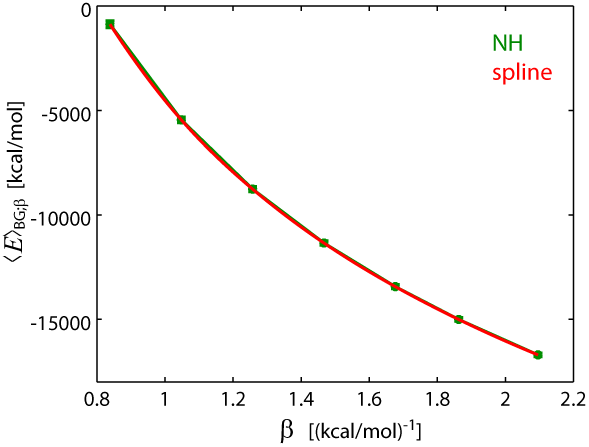
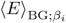 for the NH simulations of chignolin at seven temperatures (green). Their spline interpolation is shown by red.

It would be useful to note approaches other than that in Protocol (II). The simplest approach is the zeroth-order approximation of 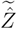 of Eq. (38) such that 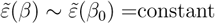 for a certain *β*_0_. Our previous study [25] can be interpreted as implicitly taking this approach (owing to the fortuitous use of the gamma function for *f*_TGT_), and suggests the difficulty in realizing a broad distribution for *β* with this approximation. As a second approach, a first-order approximation of 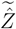 was used via a linear 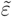 such that 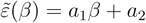, where the coefficients were determined to nicely fit the energy average curve as depicted in Fig. S3. However, this approximation failed to generate a sufficiently broad distribution of *β* to proceed to Protocol (IV), and instead was trapped at a certain *β*.

The success of the current approach in Protocol (II) can be judged by the results of the cNH simulation in Protocol (III). Figure 10(a) shows the distribution of the inverse temperature *β* obtained in Protocol (III). The region from *β*_L_ to *β*_R_, viz., the temperature range from 240 K to 600 K, was sampled. This attainment via the simple procedure is not a trivial task [25]. Discrepancies between the simulated distribution and the expected theoretical distribution [∝ Eq. (26c)] do not originate from sampling insufficiency, but from the discrepancies between the estimated 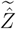 and the exact 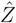. In fact, the marginal distributions of the variables *ζ, 𝒫*, and *η* were sufficiently accurate, as shown in Fig. 10(b)—(d), where the standard deviations from the theoretical distributions are 5.3 × 10^−5^, 8.0 × 10^−5^, and 5.5 × 10^−3^, respectively. Thus, we can consider that the simulated distribution of *β* agrees with the input distribution using the current 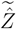, and that the approximation between *P*_B:out_ and *P*_B_ in Eq. (40) is sufficient to proceed to (IV).

**FIG. 10.**
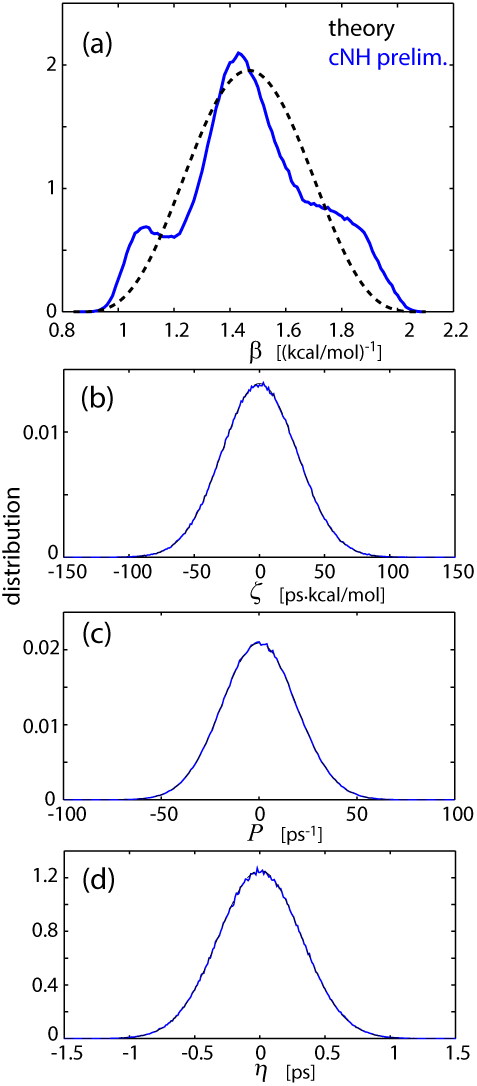
Distribution densities of (a) *β*, (b) *ζ*, (c) *𝒫*, and (d) *η* for cNH preliminary run of chignolin (blue) and theory (black).

Thus, we used the scheme provided in Sec. IV C as Protocol (IV). Figure 11 shows 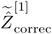, the correction of 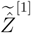 [see Eq. (43)], where the output distribution values 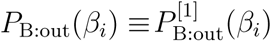 were picked up for *i* = 1, *…*, *M* ′ = 28 points to define the spline function. Note that *M* ′ was larger than *M* = 5, the number of the points taken in Protocol (I), and this choice should be applied to many cases. This is because, since the data *P*_B:out_(*β*) generated by the cNH enhanced sampling [Protocol (III)] would be statistically reliable, and since the pick-up procedure is cost-free, the use of many *P*_B:out_(*β*_*i*_) is recommended to give an accurate 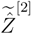 in Protocol (IV). In contrast, less accurate data ⟨*E*⟩_BG;*β*_ generated by normal BG MD [Protocol (I)] limits the use of many 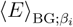 to give a tolerable 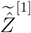 Protocol (II). For both ends *β*_0_ = *β*_L_ and *β*_*M*_*1*+1 = *β*_R_, considering the statistical insufficiency of *P*_B:out_(*β*) near these *β*_*i*_s, we set the plots to define the spline to obtain the smooth extrapolation of the curve. This is allowed since our interest is in sampling the main region of *β*, instead of the tail region.

**FIG. 11.**
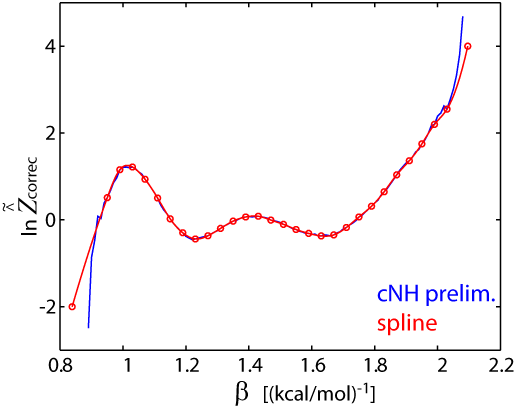
ln 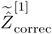 [Eq. (43)] as a function of *β* for the chignolin system.

As expected, 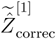 takes values of approximately 1.0 and reflects the discrepancy between the output distribution density 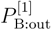 and the expected density *f*_TGT_ [see Eq. (26)], so that 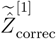 and the discrepancy have similar shape. For example, the increment of 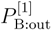 from *f*_TGT_(*β*) at *β* = 1.1, 1.4, and 1.9 observed in Fig. 10(a) clearly correspond to the increase in 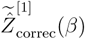 around these *β* values, respectively, observed in Fig. 11. Conversely, the decrease in the distribution at *β* = 1.2 and 1.7 corresponds to the decrease in 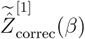 at these *β*s. Thus, the correction we should take 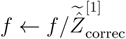 in the input of the cNH production run is expected to lead to the targeted correction of the distribution to eliminate the discrepancies.

Using the corrected (factorized-)partition function 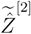 as the input of *f*, we conducted the cNH production run [Protocol (V)]. The resulting distribution of *β* obtained in this run is shown in Fig. 12(a). The distribution is more similar to the expected theoretical distribution compared with the result of the preliminary run [Fig. 10(a)]. Small discrepancies are still observed, but this result achieves our purpose, viz., to easily obtain the distribution of *β* that approximates the target distribution, as demonstrated in Sec. III. The sampling results are considered sufficient, as the distributions of *ζ, 𝒫*, and *η* are accurate [Fig. 12(b)—(d)]. The standard deviations from the theoretical distributions were 5.3 × 10^−5^, 7.3 × 10^−5^, and 6.8 × 10^−3^, respectively.

**FIG. 12.**
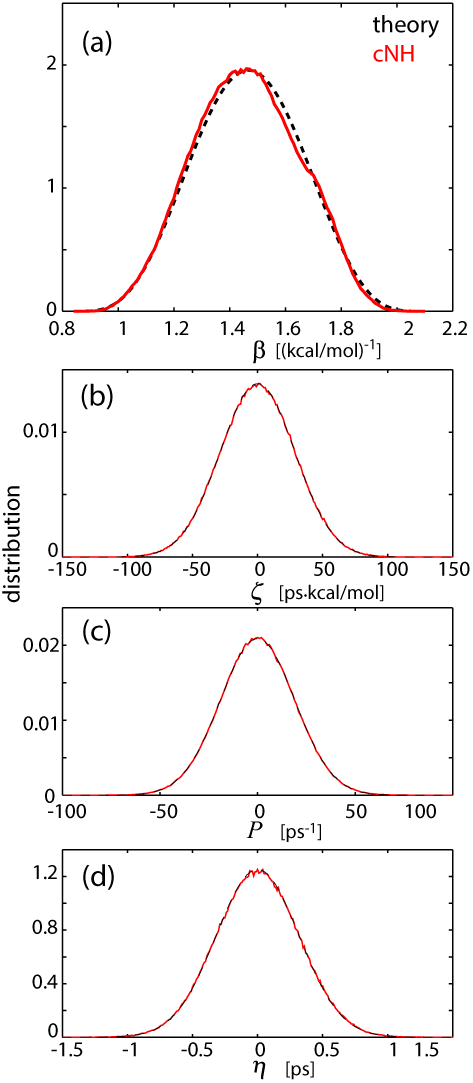
Distribution densities of (a) *β*, (b) *ζ*, (c) *𝒫*, and (d) *η* for cNH production run of chignolin (red) and theory (black).

The averages of physical quantities under the BG distribution also agreed well between the preliminary cNH run and production cNH run. See the results shown in Fig. S4 for the potential and kinetic energy distributions obtained by the reweighting to 300 K. This *a posteriori* estimate indicates that the sampling in the preliminary run was sufficiently convergent and suggests the validity of the theoretical argument (Sec. IV C) that the average of any physical-space quantity *A*(*x, p*) in the target distribution can be accurate despite 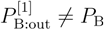. In practice, it is important that the accurate BG distribution is generated even if the output distribution density deviates from *f*_TGT_.

Figure 13 shows the TS potential energy in the (*β, ϵ*) space, as detailed in Sec. V, for this system. In contrast to the 2HO system, since the total energy *ϵ* = *E*(*x, p*) in the effective energy region Γ_E_ is negative, as observed in many biomolecular simulations, the first term *ϵβ* of the potential (44) decreases with respect to *β* [Fig. 13(a)]. On the other hand, the second term ln 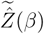 increases with respect to *β* [Fig. 13(b)]. This is because 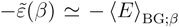 is positive and monotonically increases (recall Assumption 1 holds) in *I*_B_, so that it follows from the integration in Eq. (38). Hence, the sum of these two terms results in the valley shape (Fig. 1 actually shows these two terms) and so does the sum of all the four terms, as shown in Fig. 13(c) (note that the third and fourth TS potential terms are the same as that for the 2HO system, except for the irrelevant shift and the values of *β*_L_ and *β*_R_, and they are represented in Fig. 2). The success in making the valley shape with these two systems also indicates that the current method is robust on energy zero.

**FIG. 13.**
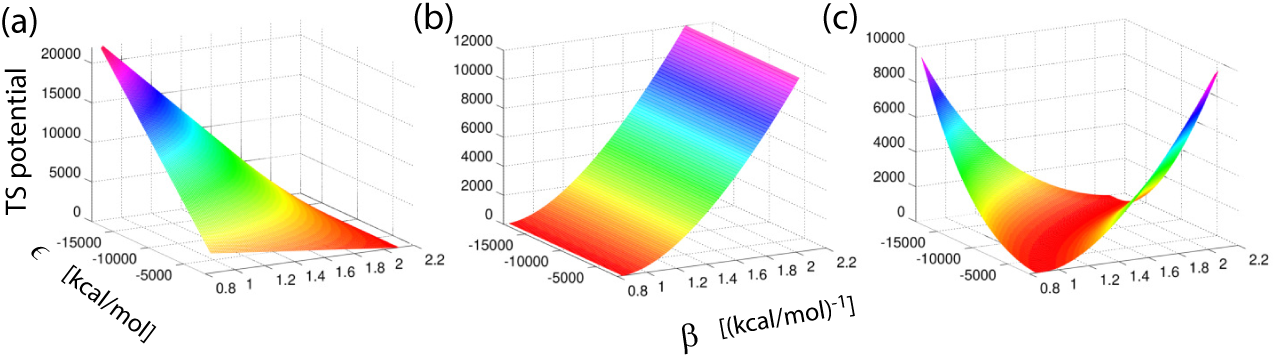
(a) *ϵβ*, (b) ln 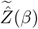, and (c) all terms of TS potential as a function of *β* and *ϵ, 𝒱*′ (*β, ϵ*), for the chignolin system. The shifting of TS potential is carried out for (a) and (c) according to Eq. (72).

Under this potential, the dynamical inverse temperature *β* should oscillate, and its motion should derive a motion of the PS total energy *ϵ* and receive its feedback. The trajectory of (*β, ϵ*) in Fig. 14(a) clearly shows that *β* and *ϵ* lie along the 2D TS potential minima. The target region *β*_L_ = 0.84 < *β* < *β*_R_ = 2.1 was attained by the trajectory in this 2D projective space. Here, as for an energy width obtained at any fixed *β*, one might think it so narrow as to contradict the ergodicity; however, it does not. The energy values varied over a wide range reaching as high as 14000 kcal/mol, and the individual widths are consistent with the probabilistic viewpoint: the probability density for which the total energy attains *E*(*x, p*) = *ϵ* under the condition that *β* =constant is, as discussed in Sec. V A [Eq. (57)], proportional to the probability density for *E*(*x, p*) = *ϵ* under the BG distribution at temperature 1*/k*_B_*β*. In fact, the individual energy widths obtained from the cNH agree with those obtained by the NH EOMs at several temperatures (= 300, 350, 400, and 450 K) in a wide range from 1*/k*_B_*β*_R_ = 240 K to 1*/k*_B_*β*_L_ = 600 K, as exhibited in Fig. 14(a). This also clearly indicates that the cNH generates many BG distributions. In addition, it was confirmed from the distributions of the PS temperature *T*_P_ = 2*K*(*p*)*/nk*_B_ and the PS potential energy *U* (*x*) shown in Fig.14 (b). Wide-range sampling in the PS is thus indicated.

**FIG. 14.**
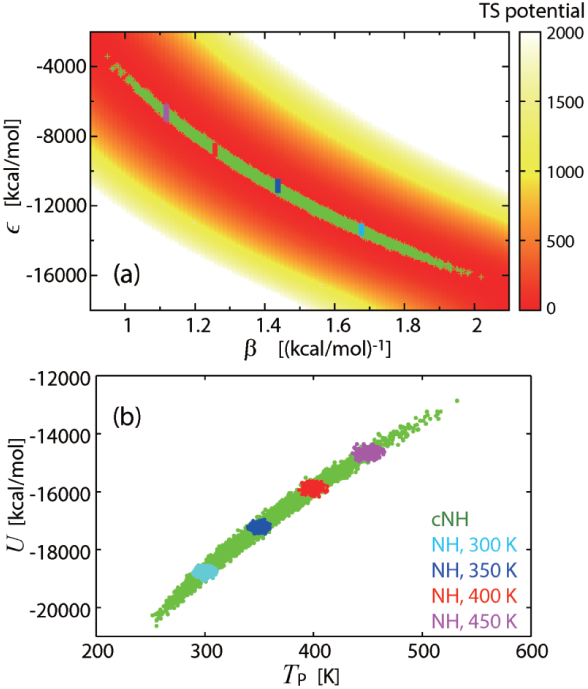
Scatter plots of (a) *β* with *ϵ* and (b) physical temperature *T*_P_ with potential energy *U* for chignolin. Shown are for cNH production run in green, and for the NH simulations at 300 K (cyan), 350 K (blue), 400 K (red) and 450 K (magenta). In (a), TS potential *𝒱*′(*β, ϵ*) is also shown, where the points for *𝒱*′(*β, ϵ*) > 2000 are omitted for clarity.

Figure 15 shows the time courses of *β* and *E* during a short period. Fundamentally, the behavior of these two quantities is similar to that observed in the 2HO. They are (negatively) correlated according to the characteristic relationship (64), and switching between processes (i) and (ii) was also observed. Terminations at *β* = *β*_L_ and *β*_R_ due to the restraint potential term can be admitted.

**FIG. 15.**
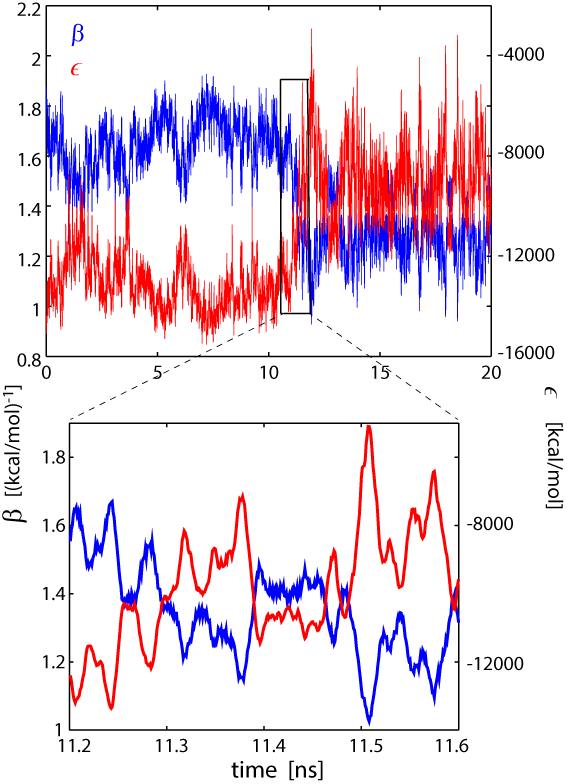
Time courses of *β* (blue) and *ϵ* (red) for cNH production run of chignolin. Close-up view is also shown.

Figures 16(a)–(d) indicate that these mechanisms are maintained for a longer period. This can be justified by the correlations between *ϵ* = *E*(*x, p*), *T*_D_ = 1*/k*_B_*β, T*_P_, and *U* (*x*), as clearly observed in the figures. The relationship between the mechanisms and these correlations is interpreted as follows. First, the process “*β* decays [grows] ⇒ *E*(*x, p*) grows [decays]” in Eq. (64) implies that “*T*_D_ grows [decays] ⇒ *K*(*p*) grows [decays]” and “*T*_D_ grows [decays] ⇒ *U* (*x*) grows [decays],” where the former seems natural. The latter may not be rigorous, but it is natural to think the grow [decay] in *K*(*p*) leads in many cases to the grow [decay] in *U* (*x*). Second, “*E*(*x, p*) grows [decays]⇒ *T*_D_ grows [decays]” is indicated via Eq. (64). Hence, the correlations of *ϵ, T*_D_, *T*_P_, and *U* (*x*) originated from the mechanisms. Note that the invariant [Eq. (109) with *T*_0_ = 300 K] was conserved in a tolerable range, indicating the success of the numerical integration. The correlation leads to enhanced sampling in the atomic conformational space as seen in Fig. 16(e), where the root mean square displacement (RMSD) of the chignolin molecular structure from the experimental data takes various values compared with that obtained in conventional BG MD at room temperature.

**FIG. 16.**
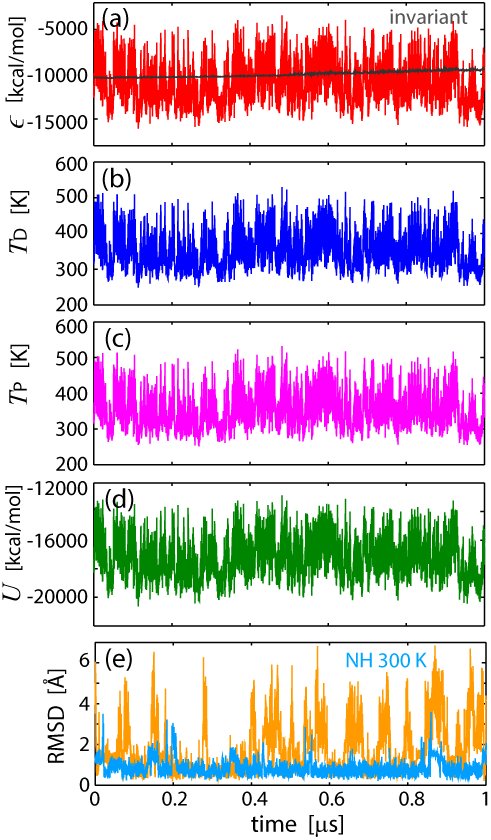
Time courses of (a) total energy *ϵ* and invariant [Eq. (108) at *T*_0_ = 300 K, gray], (b) dynamical temperature *T*_D_, (c) physical temperature *T*_P_, (d) potential energy *U*, and (e) C_*α*_-RMSD for cNH production run of chignolin. In (e) the time course for the NH simulation at 300 K is also shown as cyan.

For a specific molecular structure, the free energy landscapes spanned by two distances between hydrogen-bonding atoms, Asp3N–Thr8O (*d*_38_) and Asp3N–Gly7O (*d*_37_), in the BG distributions were calculated. The BG distribution at several temperatures were realized by the cNH with reweighting. These two hydrogen bonds are indicators of the formation of native and misfolded structures. Figure 17 (a) clearly shows the two minima, which correspond the native (right) and misfolded (left) structures, and the whole landscape including these minima. This is contrary to the scarce island [Fig. 17 (d)] obtained in conventional MD, formed mainly from the trap around the initial native structure. In fact, the reweighting result from the cNH indicated in Fig. 17 (b) implies that it should be explored more frequently between the folded and misfolded states. This transitive behavior is critical at folding temperature 350 K (the temperature at which the folded state is observed equally against unfolded states), as seen in Fig. 17 (c). Beyond this temperature, almost random motion in the accessible area was observed [Figs. 17(e) and 17(f)]. This result was unknown in our previous work [25], which had difficulty in achieving a broad temperature region for sampling. The current method can overcome this problem.

**FIG. 17.**
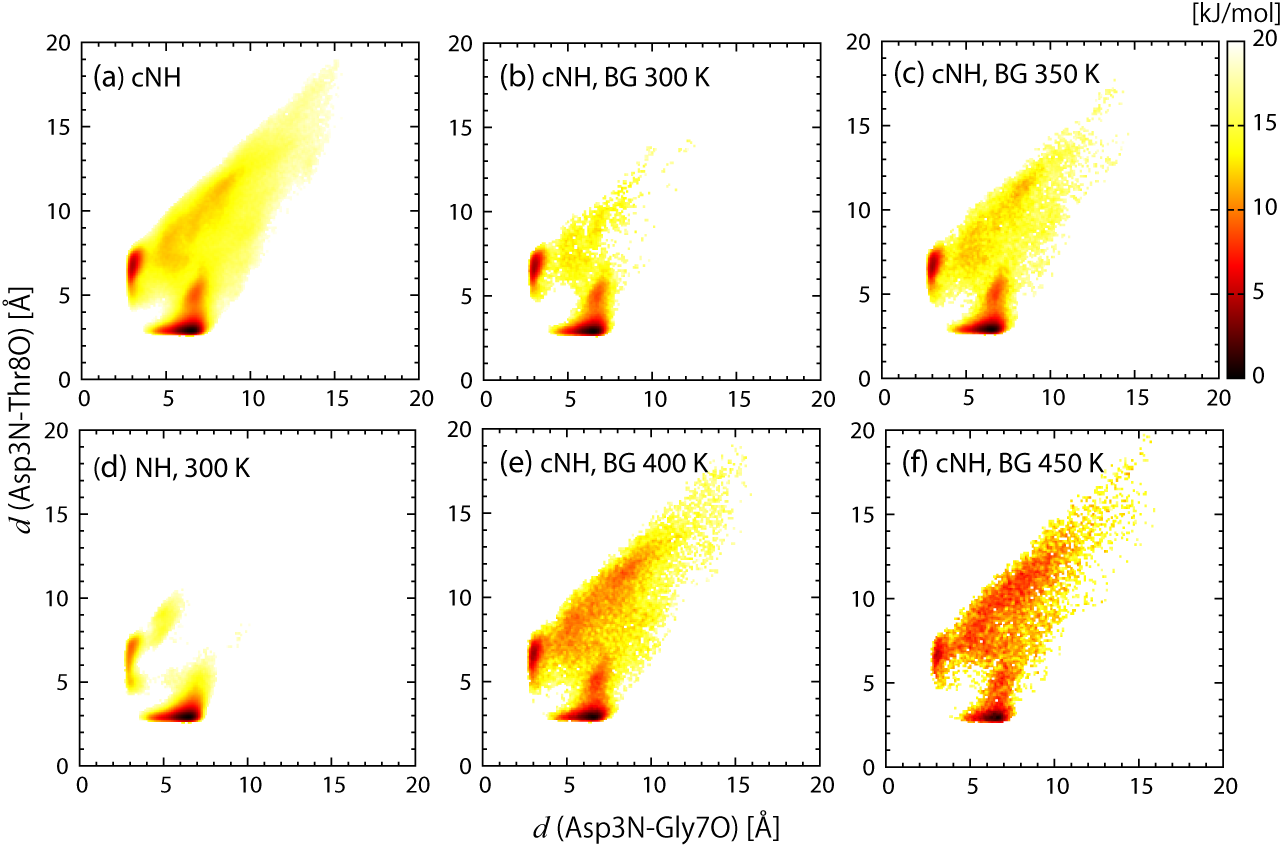
Free-energy profiles, −*k*_B_*T* ln *P*, in the plane of two hydrogen-bond distances for the chignolin system, after reweighting to the BG distribution at *T* = 300, 350, 400, and 450 K. Those of the cNH production run before reweighting (*T* = 343 K, which is the mean temperature) and the NH simulation at 300 K are also shown.

## VII. CONCLUSION

The cNH produces continuously spreading inverse-temperature distribution to the PS via deterministic and continuous time development. It uses a single replica involving the PS and TS, where the temperature (ex)change is automatically done. The current work provides a method for determining the weight of the *β* distribution to realize effective PS state sampling. This corresponds to getting sufficient information with the PS partition function in order to output a distribution that is tolerably similar to the target distribution *f*_TGT_ in a major region of *β*. For this purpose, initial data 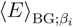 and a spline interpolation were employed, and the integration formula and simple correction were developed to complete Protocols (I)–(V), which do not require many iterations nor special reaction coordinates. The obtained knowledge is input into the potential of the TS, which is coupled to the PS in the cNH original framework.

The resulting TS potential in a 2D (*β, ϵ*)-space forms a *valley*, and the potential minimum path forms a *river* at the bottom of the valley. This path approximately becomes the (*β*, ⟨*E*⟩_BG;*β*_) curve, where ⟨*E*⟩_BG;*β*_ is the energy average under the canonical ensemble at temperature 1*/β*. Along this path, the dynamics of (*β, ϵ*) is mainly explored. Since predetermined terminal *β* values (*β*_min_ and *β*_max_) along the path can determine the terminal energy values, sampling over a wide energy range is possible. The direction of the (*β, ϵ*) movement along the river is derived from the interactions between the PS and TS. The effect of the TS on the PS specifies the direction: 1*/β* grows [decays] ⇒ *E*(*x, p*) grows [decays]. The effect of the PS on the TS specifies the reverse: *E*(*x, p*) grows [decays] ⇒ 1*/β* grows [decays]. These directions are mostly valid for *U* (*x*), as well as *E*(*x, p*). These dynamics along the 1D path in the (*β, ϵ*)-space was revealed to be the origin of the correlations between *E*(*x, p*), *U* (*x*), and 1*/β*. The specific enhanced sampling in the cNH is attained by the *mutual boost* effect: *E*(*x, p*) grows ⇒ 1*/β* grows ⇒ *E*(*x, p*) grows. This effect is characteristic but does not always hold owing to perturbations via the fluctuations, meaning the occurrence of the reverse: 1*/β* grows ⇒ *E*(*x, p*) decays. This occurrence triggers the effect in a reverse manner: *E*(*x, p*) decays ⇒ 1*/β* decays ⇒ *E*(*x, p*) decays. The termination of the *β* movement occurs at both ends *β*_min_ and *β*_max_, owing to the restraint effect of *f*_TGT_ and the Jacobian term.

To the best of our knowledge, this is the first method realizing the idea of the relationship between deterministic dynamics and (*β, E*)-space sampling. We developed a clear dynamical mechanism to explain a feature of enhanced sampling such that the temperature-space “random walk” enhances the energy-space “random walk.” The realization of the idea of the dynamics and sampling naturally arose by setting *f* under the static probability theory formalism of double density dynamics. This general formalism also enables us to extend *ρ*_E_ from the BG distribution to any distribution. These mechanisms were confirmed with numerical examples on a model system and an explicitly solvated protein system, indicating the accuracy and simplicity of the method, which are critical measures for applications. The excellent performance of the method is expected to be improved even further in the future depending on the details of the target PS and *f*_TGT_.

## ACKNOWLEDGMENTS

This work was supported by a Grant-in-Aid for Scientific Research (C) (17K05143) from JSPS and the “Development of innovative drug discovery technologies for middle-sized molecules” from Japan Agency for Medical Research and development, AMED.

## Appendix A: Spline Approximation

### Definition

In the cubic spline scheme, *S*_*i*_ (*i* ∈ *ℳ*) is a third-order polynomial defined by

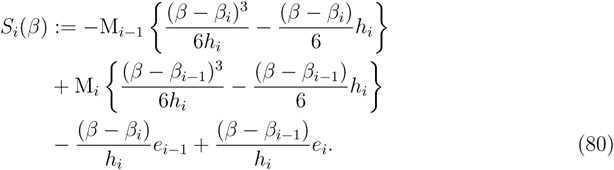

Here, M_*i*_ becomes the Hessian of *S*_*i*_ at *β* = *β*_*i*_ and they are consistently determined to yield 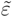 to be smooth (*C*^2^). Specifically, 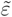 given by (37) and piecewise defined by (39) becomes *C*^2^, if {M_*i*_}_*i*=0,*…,M*+1_ satisfies

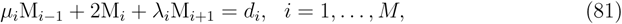

where

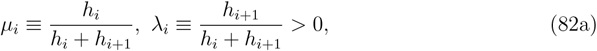

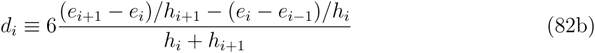

for *i* = 1, *…*, *M*. To uniquely determine *M* + 2 constants {M_*i*_}_*i*=0,*…,M*+1_, we need two conditions in addition to the *M* conditions (81). For this, we set the following endpoint conditions:

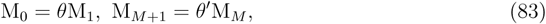

where *θ, θ*′ ≥ 0 are input parameters set to *θ, θ*′ ∼ 1 such that the behavior of 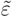 at the both ends *β*_L_ = *β*_0_ and *β*_R_ = *β*_*M*+1_ should be similar to that of neighboring points, as long as we take a sufficiently large *M*. Thus, the required condition of {M_*i*_}_*i*=0,*…,M*+1_ reads

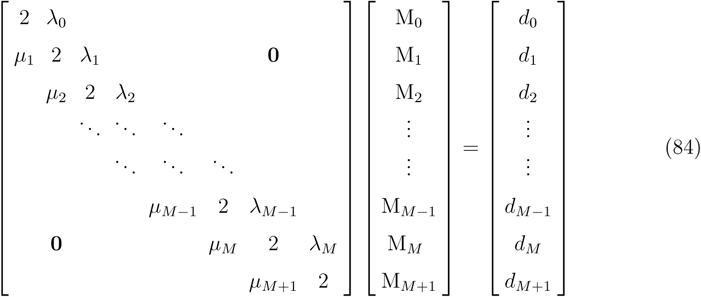

with the parameters given by Eq. (82) and

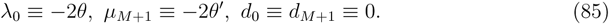

By solving linear Eq. (84) (defined by the matrix with adjustable parameters *θ* and *θ*′), we have the required {M_*i*_}, and thus have *C*^2^ function 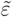 [defined by Eqs. (37), (39), and (80)].

### Quantities

We give specific forms of quantities used in the current method. First, we need − ln 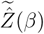 for *β* ∈ *I*_B_, which is utilized in TS potential (28) and appears in the invariant function [Eq. (108)]. The integration in (38) yields

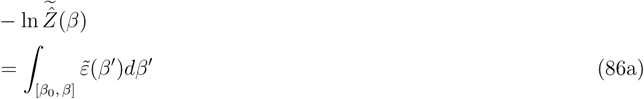

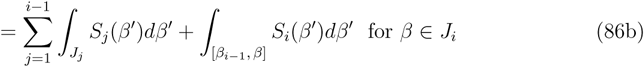

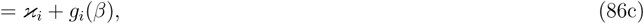

where *g*_*i*_(*β*) and ***x***_*i*_ are a polynomial and a constant, respectively, defined by

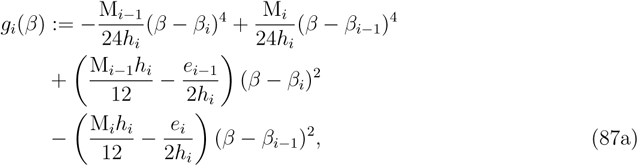

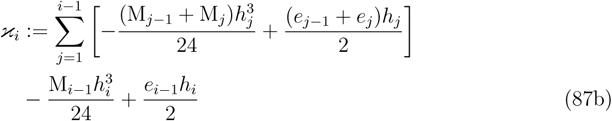

for *i* ∈ *ℳ*. Here, {M_*i*_} are obtained as a solution of Eq. (84).

Second, we need 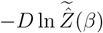, which is used in the TS force [see Eq. (46)] and easily obtained for any *β* ∈ *I*_B_ as

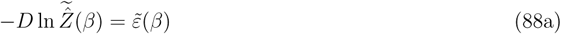

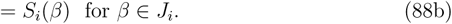

The mapping from *β* ∈ *J*_*i*_ to the index *i* for implementing Eqs. (88b) and (86c) can be easily constructed.

Last, we need *ρ*_RE_(*E*(*x, p*)) defined by Eq. (17) for the reweighting via Eq. (18). We have

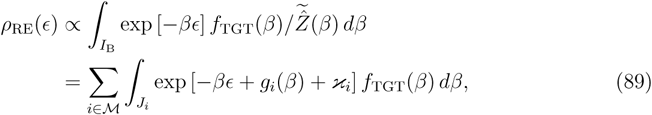

which can be evaluated numerically for each *ϵ* (normalization constant is not needed for our purpose). In practice, *ρ*_RE_(*ϵ*_*i*_) is evaluated at discrete points *ϵ*_*i*_, and a spline-function interpolation can be used to define *ρ*_RE_(*ϵ*) for any *ϵ* in a similar manner.

## Appendix B: General case of the TS potential

Changing *ρ*_E_ from the BG case [Eq. (8)] to a general case yields the change in the major part of the TS potential, viz., from Eq. (47) into

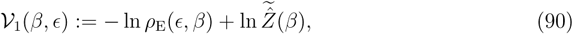

yielding the TS force

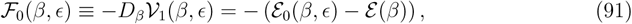

where

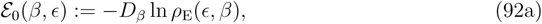

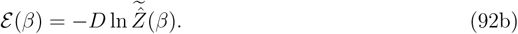

Since *ε* can be approximated by Eq. (30) such that

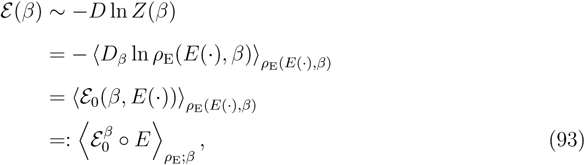

the TS force maintains the structure described by the difference between an energetic quantity and its PS-space average:

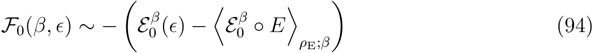

or

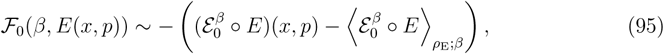

which is analogous to Eq. (63), where *E* is generalized into 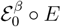 (note 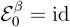 in the BG case) and 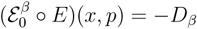 ln *ρ*_E_(*E*(*x, p*), *β*) has a dimension of energy.

For each *ϵ* = *E*(*x, p*) ∈ Γ_E_, the stationary point *β* is characterized by

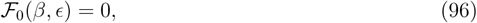

which can be locally solved with respect to *β*, following the implicit function theorem. We simply assume that it can be solved globally such that

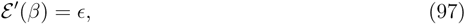

Implying

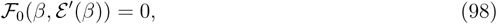

for any *β* ∈ *I*_B_. We then suppose that *ε*′ is injective and Γ_E_ ⊂ *ε*′(*I*_B_), and thus have the set of stationary points

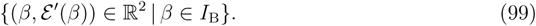

Eqs. (52) and (55) read

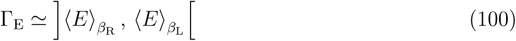

and

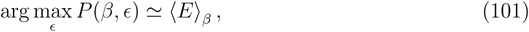

respectively, where

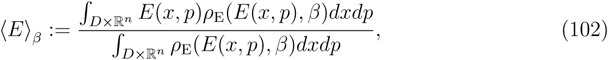

and Eq. (101) can be satisfied for *ρ*_E_ having a nice property. To maintain a parallel discussion of Sec. V A, we further require additional conditions:

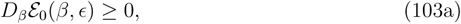

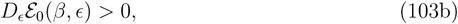

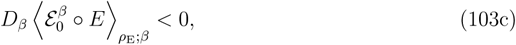

for any (*β, ϵ*) ∈ *I*_B_ × Γ_E_. These are conditions on *ρ*_E_ and satisfied with the BG case; hence, they are used to set *ρ*_E_ in the general case. Then, Eqs. (103a) and (103c) lead to the concavity of the TS potential,

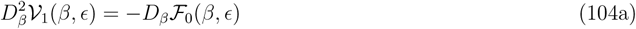

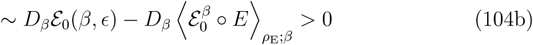

for any (*β, ϵ*), implying that *𝒱*_1_ is a “valley” and subset (99) is an underlying “river.” Note that Eq. (104b) can be replaced with Eqs. (103a) and (103c) as a weaker condition. As a generalization of Eq. (64), a mechanism

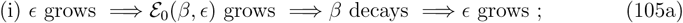

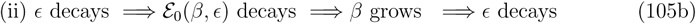

now holds, where the first “⇒” relation is derived from Eq. (103b), the second “⇒” relation is supported by Eq. (95), and the third “⇒” relation is suggested by the same reason discussed in section V A. The switching between process (i) and process (ii) should be made by the allowance for the perturbation of the fluctuations in the third “⇒” relation, as stated in Sec. V B, and by the addition of the other terms ℱ(*β*) into *𝒱*_1_(*β, E*).

## Appendix C: Integrator of the EOM

For numerical integration, a target ODE, 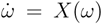, defined on a phase space Ω is Eq. (4). Here, we focus on the case in which *ρ*_E_ is described by Eq. (8) with *m* = 1 and *c*_*T*_ = 0 in Eq. (7a), for simplicity. Consult Ref. [52] for a general case. The extended space formalism [40, 41] defines an extended ODE on an extended space Ω′ ≡ Ω × ℝ by

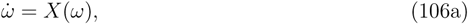

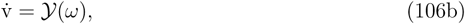

where v is introduced as a variable on the additional space ℝ, and

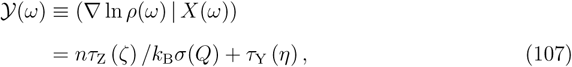

with *ρ* given by Eq. (6). We then have an invariant *L* for ODE (106), i.e., *L*(*ω*′(*t*)) is constant for any time *t* for an arbitrary solution, *t* ↦ *ω*′(*t*) ≡ (*ω*(*t*), v(*t*)), of ODE (106). The invariant function is given by

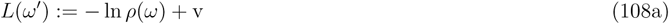

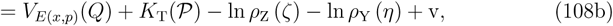

where *V*_*E*(*x,p*)_(*Q*) is given by Eq. (28). Equation (106) does not destroy the solutions of the originally given ODE (106a). Monitoring the value of Eq. (108) enables us to check the numerical error to integrate ODE (106). It is convenient to use an energy analogue

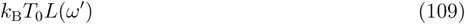

with a certain temperature *T*_0_, instead of the nondimensional *L*(*ω*′), to observe the numerical error compared with the fluctuations of the PS energy. In practice, the initial value of v is set to be 0.

An explicit, symmetric, second-order integrator with time step *h* for ODE (106) [31] can be given by

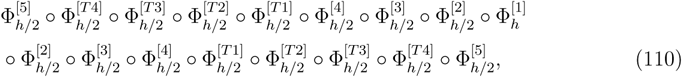

where the maps on Ω′ Are

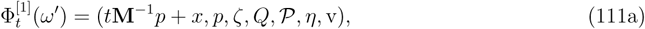

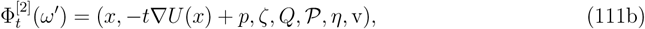

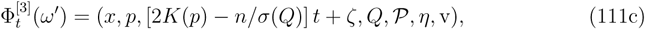

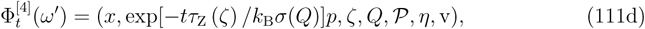

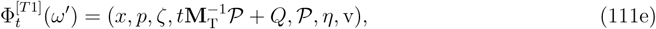

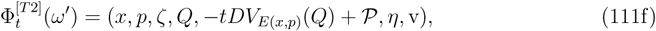

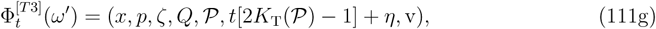

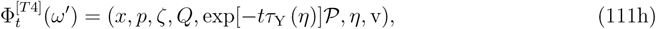

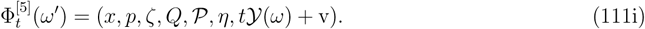

The part that is far more time consuming than other parts is the estimation of the atomic force −∇*U* (*x*) ∈ ℝ^*n*^, for which the one-time evaluation suffices in Eq. (110).

